# Improved metabolic syndrome and timing of weight loss is crucial for influenza vaccine-induced immunity in obese mice

**DOI:** 10.1101/2022.04.16.488487

**Authors:** Rebekah Honce, Ana Vazquez-Pagan, Brandi Livingston, Sean Cherry, Virginia Hargest, Bridgett Sharp, Lee-Ann Van de Velde, R. Chris Skinner, Paul G. Thomas, Stacey Schultz-Cherry

## Abstract

Persons with obesity are at higher risk for developing severe complications upon influenza virus infection making vaccination a priority. Yet, studies demonstrate vaccine responses are less effective in obese hosts. In these studies, we examined how the timing of weight loss influenced influenza vaccine efficacy in male and female diet-induced obese mice. Here, we show weight loss post-vaccination is insufficient to rescue poor vaccine efficacy; however, weight loss occurring pre-vaccination successfully improves outcomes at viral challenge. Pre-vaccination weight loss improved vaccine immunogenicity and restored a functional recall response at challenge. Through tracking sera metabolic biomarkers, we propose the metabolic state at the time of vaccination is predictive of vaccine immunogenicity. Altogether, these findings highlight how timing of host-directed interventions is vital when seeking to improve influenza vaccine immunogenicity in obese hosts.

## Introduction

Obesity is a leading cause of global morbidity as it underlies many non-communicable diseases, such as cardiovascular disease and several cancers^1^. As obesity’s prevalence increases, we have begun to appreciate its impact on susceptibility to and severity of communicable diseases^2,3^. Epidemiological and empirical studies suggest obesity increases severity of bacterial^4–6^, viral^7^, fungal^8^, and parasitic^9,10^ infections. After the 2009 H1N1 influenza virus pandemic, obesity was identified as an independent risk factor for hospitalization, need for intensive care, and death upon infection^11^. These trends are mirrored in the COVID-19 pandemic^12^.

Prevention through vaccination is the best control strategy for many infectious diseases. However, previous studies in our lab and others determined obesity may weaken influenza immunization efficacy^13–15^. Even with higher antigen dose and adjuvants, inactivated vaccines do not improve survival in obese mice upon viral challenge^16,17^. Vaccinated adults with obesity are twice as likely to develop PCR-confirmed influenza than people with a body mass index (BMI) of 25 kg/m^2^ or less^18^. It has been posited the metabolic consequences of obesity attenuate generation of protective immune responses post-vaccination^14,19^. Previous studies have interrogated the contribution of diet to immunity, and how flux in metabolic state can impact overall health^20,21^. In these studies, we asked if host interventions, such as weight loss, could improve immune responses to whole, inactivated influenza viruses in a mouse model of diet-induced obesity.

To answer our overarching question, we maintained wild-type C57Bl/6J mice on either high-fat, calorically dense diet (HFD) or standard chow for 16 weeks. After four months of exposure to primary diets, mice were vaccinated with inactivated H1N1 influenza virus. Following immunization, a subset of HFD-exposed mice was swapped to standard chow. This secondary diet continued for either 4 weeks (short-term) or 12 weeks (long-term), at which time homotypic viral challenge with a lethal dose of H1N1 virus occurred. Neither short- nor long-term weight loss post-vaccination improved viral clearance, seroconversion, or survival upon challenge in formerly obese mice. However, survival outcomes were improved—and indistinguishable from always lean mice—if weight loss occurred prior to vaccination. By monitoring markers of adipose tissue function, we found that high leptin, low adiponectin, and a low adiponectin:leptin ratio at the time of vaccination is associated with poor vaccine efficacy. Immunologically, metabolic dysfunction stemming from HFD exposure at the time of vaccination blunted the primary generation of CD4^+^ and CD8^+^ T cells, which remained impaired upon post-vaccination weight loss. Switching formerly obese mice to the lean control diet before vaccination resulted in gradual weight loss and return of these metabolic biomarkers to the lean baseline, restored humoral- and cellular-mediated protection at challenge and led to 100% survival in formerly obese mice. These studies suggest nutritional status at the time of vaccination fixes immunization efficacy, possibly through crosstalk between metabolism and the immune system.

## Results

### Neither short- nor long-term weight loss post-vaccination improves survival outcomes

Poor infection outcomes, regardless of vaccination status, is a hallmark of influenza pathogenesis in obese mice^16,22^. To determine if weight loss is sufficient to improve survival outcomes after immunization, we randomized whole cages to either HFD or standard chow for 16 weeks^23^ after which mice were vaccinated with whole, inactivated H1N1 influenza virus (Fig. 1a). Each experimental group consisted of n=20 mice with half serving as unvaccinated controls. At the time of vaccination, HFD and chow-fed groups had significant differences in weight (p<0.0001, Fig. 1b, c). Two weeks post-vaccination, diets were switched to create three cohorts: always obese (HFD>HFD), always lean (Lean>Lean), and formerly obese (HFD>Lean) mice. During the post-diet switch, formerly obese mice displayed notable weight loss compared to always obese mice (p<0.0001, Fig. 1b, c).

**Figure 1.**
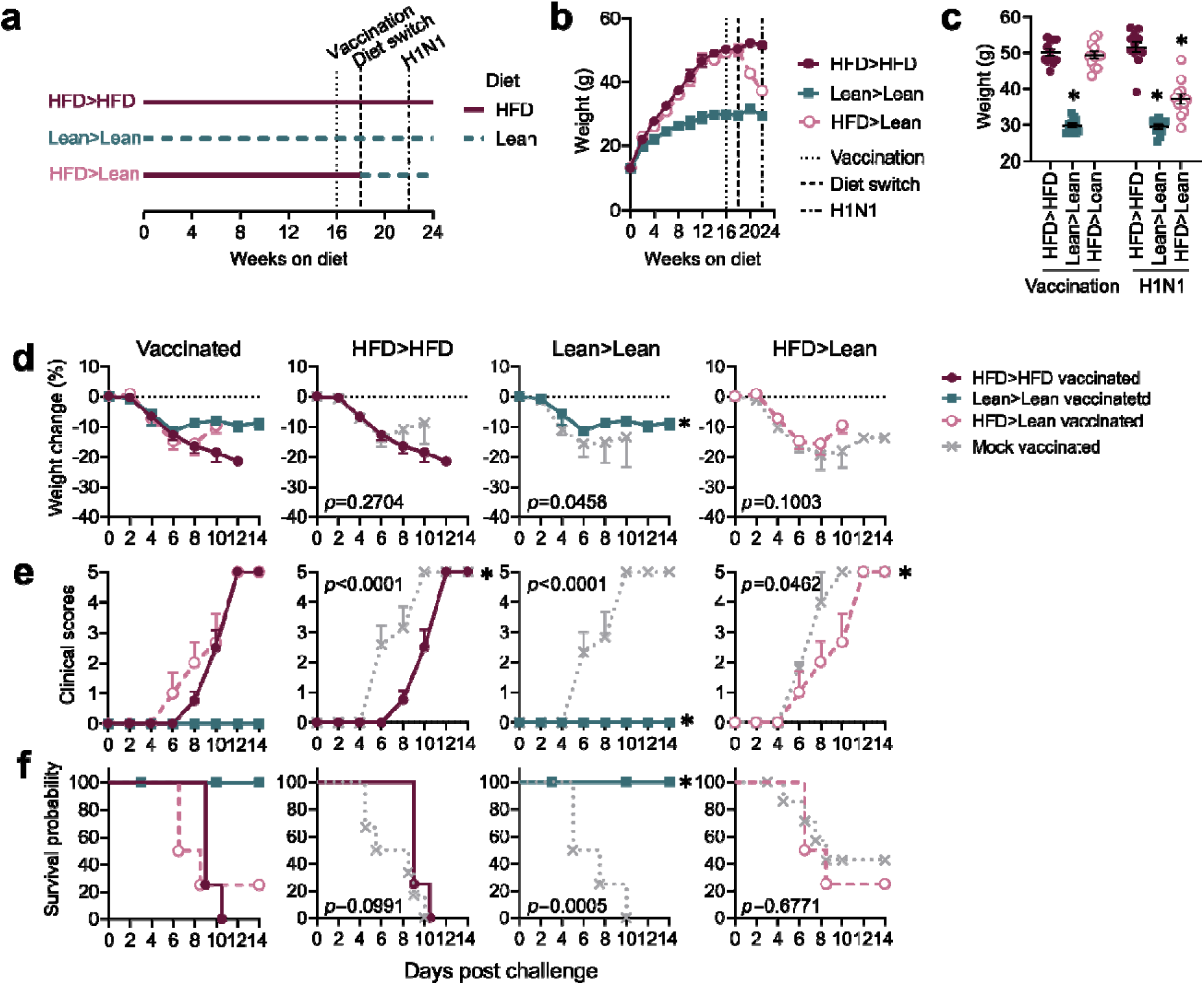
Short-term weight loss does not improve survival upon viral challenge. (a) Timeline of diet administration, vaccination, and challenge for 4-week short-term diet protocol. (b) Weights of mice through short-term diet protocol and at (b) vaccination and challenge. (d-f) Morbidity and mortality in vaccinated mice compared to mock vaccinated mice with n=10 mice/group. (d) Weight curves post-challenge with statistical comparisons made via mixed-effects model. (e) Clinical scores post-challenge with statistical comparisons made using a two-way ANOVA. (f) Survival post-challenge with statistical comparisons made using Mantel-Cox log-rank analysis. Data in (b-e) are representative of two independent experiments and are graphed as means ± standard error and in (f) as surviving proportions with censored or event animals indicated by their respective symbols. Statistical comparisons test between HFD>HFD and indicated diet group (n=10 mice/group) in (b, c) and between mock and vaccinated animals within a single diet group in (d-f). Red closed circles = always obese mice, green closed squares = always lean mice, pink open circles = formerly obese mice, and grey dashed x = mock vaccinated mice

At 1-month post-diet switch, mice were challenged with 10X mouse lethal dose-50 (MLD_50_) A/California/04/2009 (CA/09) as indicated (Fig. 1a) and monitored for morbidity and mortality. In contrast to mock- or always lean groups, vaccinated always obese mice were not protected from morbidity or mortality as 100% lethality was observed (Fig. 1d-f). This was also true in formerly obese mice. Weight loss post-vaccination failed to attenuate infection-related weight loss (p=0.1003, Fig. 1d) but did statistically reduce signs of infection (*p*=0.0462, Fig. 1e), although this is not biologically relevant as survival of formerly obese mice was not improved compared to survival rates in always lean mice (p=0.6771, Fig. 1f). Overall, vaccination resulted in survival of 100% of always lean, 0% of always obese and 24% of formerly obese mice.

We considered that longer-term weight loss may be needed to protect formerly obese mice. However, even at 12 weeks post-diet switch, vaccination did not improve survival outcomes in formerly obese mice (Supplemental Fig. 1). Weight trends were comparable to short-term diet switch mice (Supplemental Fig. 1a-c). At challenge, improvements in morbidity were noted but survival was not improved for mice vaccinated while obese (Supplemental Fig. 1d-f). Compared to mock vaccinated controls, vaccinated always lean mice were 100% protected from viral challenge (*p=*0.0001), but we observed poor survival outcomes (0% and 28%, respectively) in always obese and formerly obese mice (Supplemental Fig. 1f). In conjunction with the short-term experiment, these studies attest diet group at time of vaccination is predictive of survival at challenge in mice.

### Weight gain modestly attenuates vaccine efficacy but overall preserves survival outcomes

To determine if diet at the time of vaccination is predictive of survival at viral challenge, we next questioned if post-vaccination weight gain would diminish vaccine efficacy in lean mice. Thus, mice were maintained on diet schedules as shown in Figure 1, with a fourth Lean>HFD group added (Supplemental Fig. 2a). Post-diet switch, weight gain in formerly lean mice was robust and rapid but did not reach the same mass as mice maintained on HFD even after 12 weeks (Supplemental Fig. 2b, c). Regardless of length of weight gain, mice that were lean at time of vaccination were significantly protected from mortality upon challenge (Supplemental Fig. 2d, e), while morbidity was more affected from the diet switch, especially longer-term weight gain (Supplemental Fig. 2d, e). Formerly lean mice did show modest reductions in overall survival with short-term (75% survival) and long-term weight gain (85.7%) compared to always lean mice (100%) suggesting a small influence of diet at the time of challenge. Overall, these data suggest diet group and associated body mass at the time of vaccination influences vaccine efficacy, but diet at challenge may contribute to overall survival trends.

### No observed sex bias in diet, vaccination, or challenge outcomes

Sex biases in responses to vaccination and infection are well-established^24–26^. We have previously observed differing influenza pathogenesis and immune responses in obese male and female mice^27^, and there is a complex web of interplay among sex hormones, obesity, and immunity^28^. To examine potential sex bias in our system, female mice were subjected to the same diet and challenge protocols as described (Supplemental Fig. 3a). Growth curves and weight at vaccination and challenge were concordant with male mice, albeit females had lower overall mass^29^ (Supplemental Fig. 3b, c). Vaccination reduced morbidity in always lean and formerly obese mice (Supplemental Fig. 3d, e).

However, survival outcomes were again only improved in mice vaccinated when lean, including both always lean and formerly lean diet groups (Supplemental Fig. 3f). These findings emphasize weight loss post-vaccination is insufficient to increase vaccine efficacy in male and female mice. Further illustrating how diet group at time of vaccination is predictive of survival post-challenge regardless of weight loss or gain post-vaccination.

### Weight loss does not improve viral clearance but recovers interferon-mediated control of viral spread

To better understand why the vaccine failed to protect formerly obese mice, we monitored viral titers and spread, lung histopathology, and interferon responses. Survival trends were similar between short- and long-term diet protocols, thus, the remaining studies were performed using the short-term protocol. At days 3 and 7 post-infection (p.i.), reduced viral titers were only observed in the always lean vaccinated diet group (Fig. 2a). There were no differences between the obese or formerly obese groups. As obesity is associated with increased viral spread in the lungs^27,30^, we performed *in situ* hybridization (ISH) for the nonstructural gene (NS1, Fig. 2b). In always obese mice, marked viral spread was noted at day 3 p.i. (white arrow, Fig. 2b), while spread was limited in always lean and formerly obese mice suggesting diet group at challenge may correlate with viral spread.

**Figure 2.**
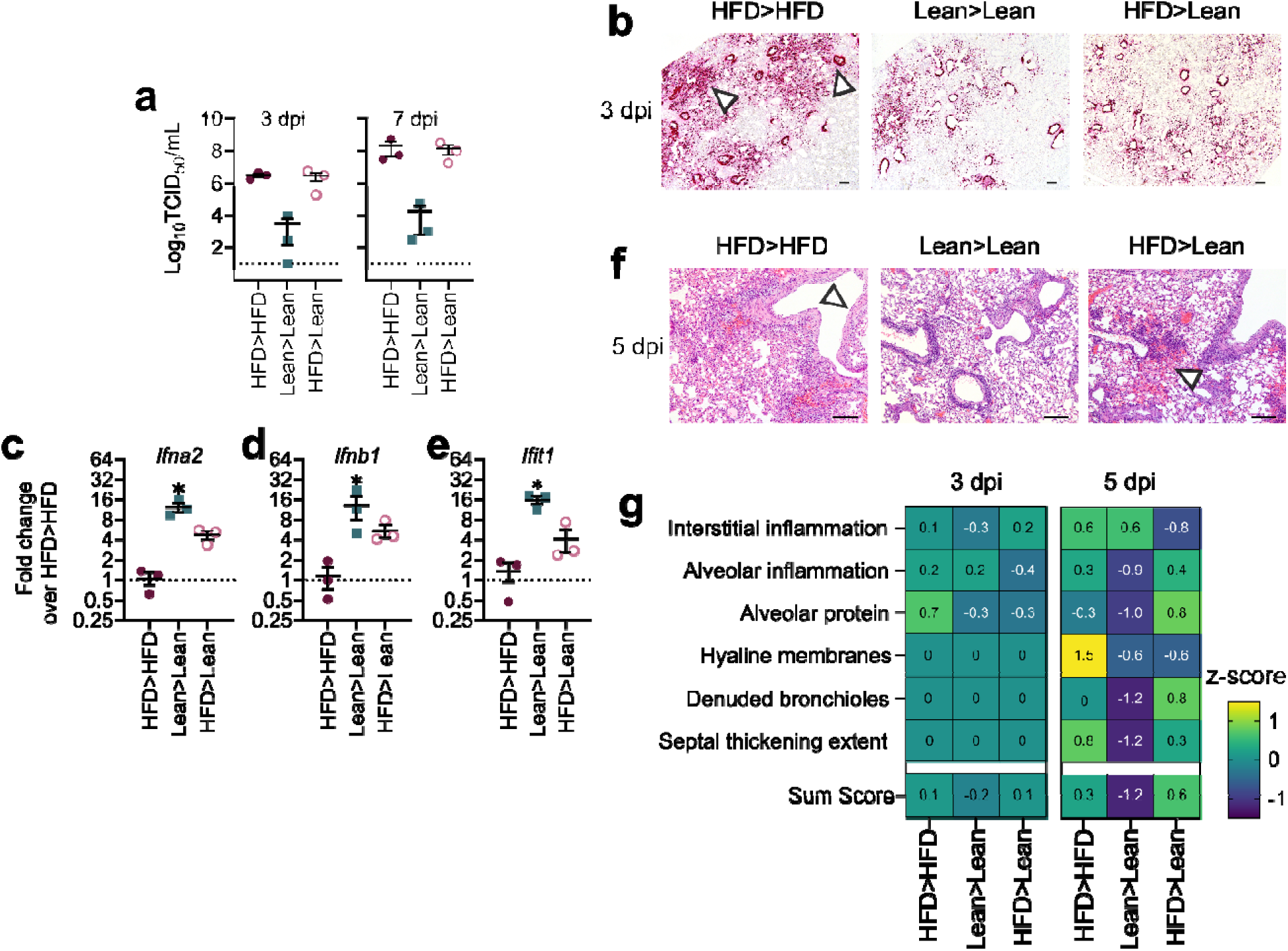
Heightened markers of inflammation despite improved viral spread in formerly obese mice. (a) Lung viral load in short-term diet switch mice at day 3 or day 7 post-challenge. (b) Lung sections at day 3 post-challenge probed for H1N1 NS1 protein through *in situ* hybridization. (c-e) Expression of (c) *Ifna2*, (d) *Ifnb1,* (e) *Ifit1* at day 3 post-challenge. Results include n=3 mice per diet per treatment and are representative of two independent experiments. (f) Representative hematoxylin and eosin-stained slides from indicated mice at day 5 post-challenge with H1N1 virus after short-term diet switch. (g) Quantification of pathological findings standardized via z- scores. Z-scores reported are an average of n=2 or 3 mice per diet group per time point. Graphed in (a, c-e) as means ± standard error and in (g) as z-scores with comparisons made between diet groups to levels in always obese (HFD>HFD) mice via ordinary one- way ANOVA with Dunnett’s correction. Red closed circles = always obese mice, green closed squares = always lean mice, and pink open circles = formerly obese mice. Scale bar equal to 4 mm.

At these early time points, viral spread is largely controlled by innate immune responses^31^. Prior studies determined interferon is sensitive to diet group, with blunted and delayed upregulation of interferon and interferon-stimulated genes evident in obese mice and obese-derived human epithelial cell cultures, possibly permitting greater viral spread^30,32^. Compared to always obese mice, always lean mice expressed higher lung *Ifna2* (*p*=0.0010), *Ifnb1* (p=0.0531), and *Ifit1* (p=0.0013) levels at day 3 p.i. (Fig. 2c-e). Formerly obese mice had a slight increase in *Ifna2* (p=0.1218) expression, suggesting a modest recovery in interferon responses. These data align with the observed differences in viral spread, reinforcing that diet group at viral challenge—but not at vaccination—correlates with interferon-mediated control of viral spread.

We next examined lung histopathology to determine if formerly obese mice, which had higher interferon levels and reduced viral spread, showed less severe inflammation upon challenge (Fig. 2f-g). Instead, vaccinated obese and formerly obese mice had more extensive lung pathology as evidenced by bronchiolar epithelium loss, interstitial edema, and alveolar fibrin/hemorrhage, with only minimal perivascular lymphoid cell infiltrates compared to always lean mice (white arrows, Fig. 2f, quantification Fig. 2g). In always lean mice, the bronchiolar epithelium was largely preserved (although flattened), interstitial edema and alveolar fibrin/hemorrhage were minimal, and perivascular lymphoid cell infiltrates were prominent. Overall, these data highlight how diet at both immunization and challenge impacts viral spread and lung architecture after challenge and contribute to overall survival.

### Vaccination while obese leads to blunted seroconversion

To assess antibody responses, hemagglutination inhibition (HAI)^33^, plus total and H1N1- specific IgG titers were quantified by ELISA at 2 weeks post-vaccination, pre-challenge (either 1 month or 3 months post-diet switch for short and long-term diet protocols, respectively) and post-challenge. Neither total nor viral-specific IgG titers differed by diet (Fig. 3a, b), which contrasts with HAI titers where only the vaccinated, always lean mice seroconverted (Fig. 3c, d). Mice vaccinated while obese did not seroconvert prior to viral challenge (Fig. 3c, d); however, post-challenge any surviving formerly obese mouse did seroconvert (Fig. 3c, d). Female mice had comparable results (Supplemental Fig. 3g).

**Figure 3.**
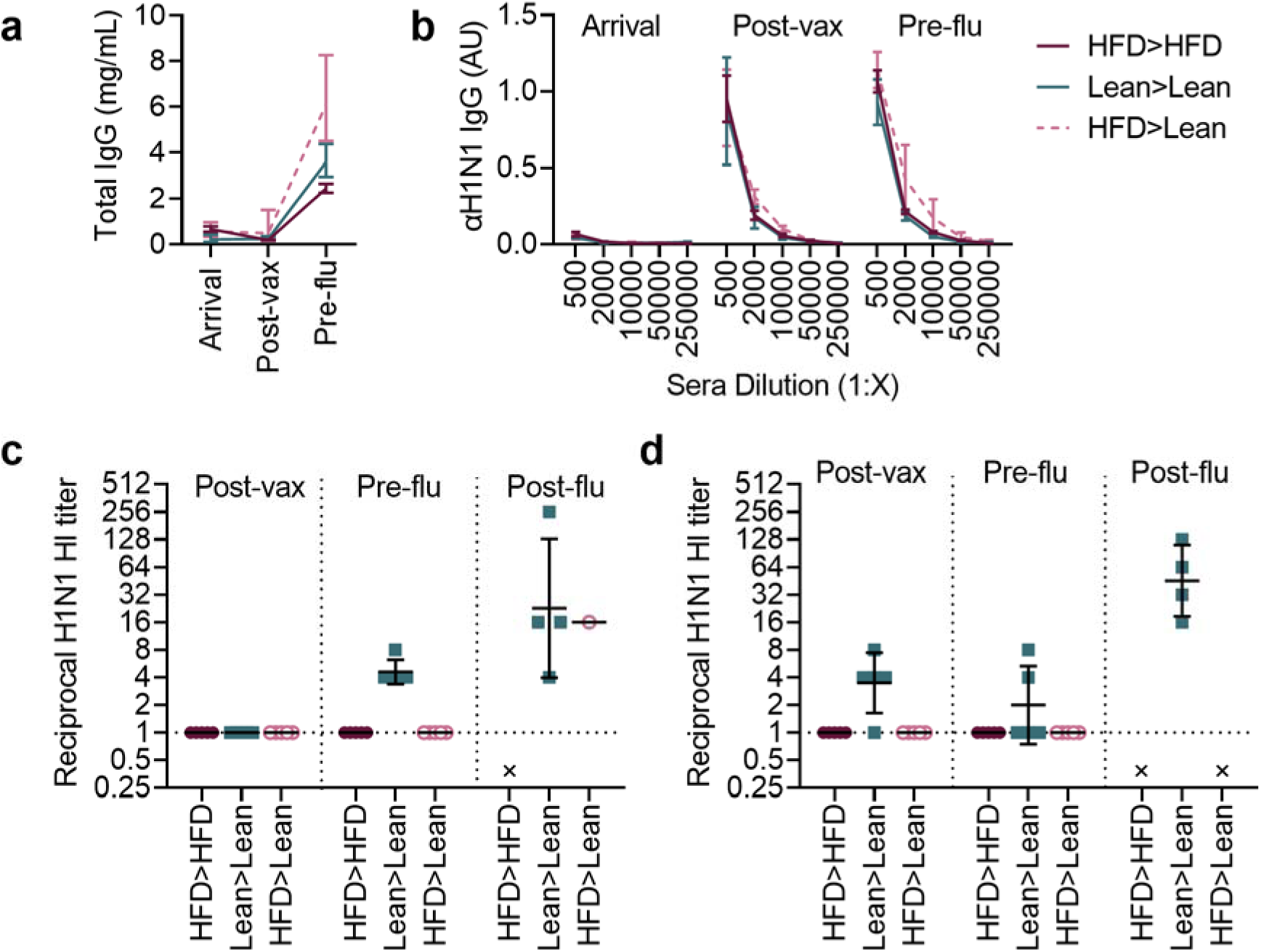
Blunted seroconversion in mice vaccinated when on high-fat diet. Total (a,b) and effector antibody levels in sera collected and quantified via (a) total and (b) viral-specific ELISA and hemagglutination inhibition (HAI) in short-term (a-c) and long-term (d) diet switch mice. Mice assayed longitudinally with n=5 per group for (c, d) and n=3 per group for (a, b) except for the day 21 post-influenza challenge timepoint in which only surviving mice were assayed. Data is representative of four independent experiments and displayed as geometric means ± standard deviation with x indicating no mice surviving to time point. Red closed circles = always obese mice, green closed squares = always lean mice, and pink open circles = formerly obese mice.

These data imply that weight loss after vaccination does not improve production of anti-influenza specific antibodies^16^.

### Blunted memory responses in vaccinated obese mice

Obese mice have an impaired memory T-cell recall response upon secondary challenge that is not reversed with weight loss^34^, potentially contributing to the poor viral clearance and survival outcomes in this model^35^. In our study, at day 7 p.i., formerly obese mice have an increasing trend in overall CD4^+^ and CD8^+^ T-cell numbers within the lung tissue, including T_EFM_ and T_CM_ cell populations, as compared to the other groups (Fig. 4a). In contrast, CD8^+^ T_EFM,_ and CD8^+^ T_CM_ numbers were comparable across all groups (Fig. 4a, Supplemental Fig. 6). Although absolute numbers differed, frequency did not (Supplemental Fig. 6), which may reflect the inherent differences in baseline inflammation and cellularity in obese mice and formerly obese mice.

**Figure 4.**
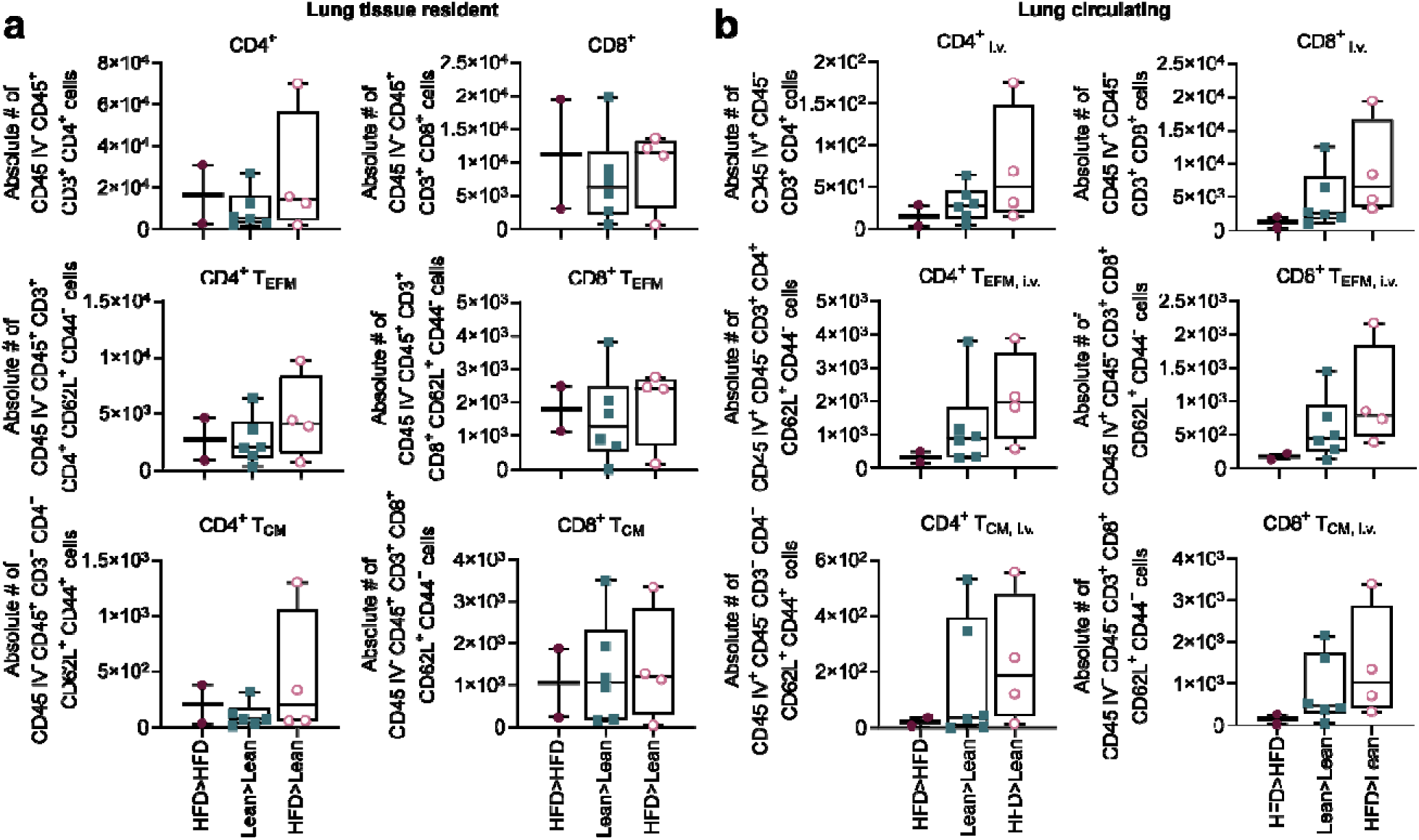
Blunted memory responses in mice vaccinated when on high-fat diet. Lungs were harvested and homogenized into single-cell suspensions and 1 × 10^6^ cells were stained for CD4^+^ and CD8^+^ T cells at 7 days post-challenge and quantified via flow-cytometry. (a) Absolute number of lung resident CD45_i.v.−_, CD4^+^ and CD45_i.v.−_, CD8^+^ T cells; CD45_i.v.−_, CD4^+^ effector memory (T_EFM_) and CD45_i.v.−_, CD8^+^ T_EFM_ cells; CD45_i.v.−_, CD4^+^ central memory (T_CM_) and CD45_i.v.−_, CD8+ T_CM_ cells. (b) Absolute number of circulating CD45_i.v.−_, CD4^+^ and CD45_i.v.−_, CD8^+^ T cells; CD45_i.v.−_, CD4^+^ effector memory (T_EFM_) and CD45_i.v.−_, CD8^+^ T_EFM_ cells; CD45_i.v.−_, CD4^+^ central memory (T_CM_) and CD45_i.v.−_, CD8+ T_CM_ cells. Data were analyzed using FlowJo version 10.8.1. Outliers in the dataset were removed using the robust regression and outlier removal (ROUT) method (Q=1%). Data include always obese (HFD>HFD), always lean (Lean>Lean), and formerly obese (HFD>Lean) mice (n = 2, n=6, n=4 for each group, respectively) and graphed as means ± standard error with comparisons made between always lean and formerly obese groups via a paired T-test. Red closed circles = always obese mice, green closed squares = always lean mice, and pink open circles = formerly obese mice.

To further distinguish circulating T-cell populations (CD45_i.v.+_, Fig. 4b), we performed an intravenous (i.v.) injection of CD45 antibody prior to lung tissue collection (Supplemental Fig. 5)^47–50^. Always lean and formerly obese mice had increased infiltration of both CD4^+^ and CD8^+^ T-cells in the lung tissue, including T_EFM,_ and T_CM_ cell populations, as compared to the always obese mice (Fig. 4b). Overall, these data suggest that, while weight loss does increase the number of lung memory T cell populations and moderately improves survival rates in formerly obese mice, it is still not sufficient to overcome the ramifications of vaccination while obese. These findings also led us to speculate that antibody-mediated immunity, and not solely a T-cell recall response, is critical for viral clearance in our model since only always lean mice, which had anti-influenza specific antibodies (Fig.3c, d) were able to clear the virus at this time point (Fig. 3a) and result in 100% survival (Fig.1f, Supplemental Fig.1f).

### Metabolic dysregulation at time of vaccination is predictive of poor primary vaccine responses

The systemic impact of obesity likely attenuates the vaccine response to numerous pathogens^13,36,37^. We observed many components of metabolic syndrome (MetS) in our obese mice, including heightened liver mass (p<0.0001, Fig. 5a, b) and lipid accumulation (p<0.0001, white arrows Fig. 5a, quantification Fig. 5c), triglyceride content (p=0.0262, Fig. 5d) and elevated oxidative markers (p=0.0393, Fig. 5e)^38,39^ compared to always lean mice. After 12 weeks of weight loss, formerly obese mice had significantly reduced liver adiposity (p<0.0001) and modest decreases in oxidative stress (p=0.1950) compared to always obese mice. These intractable systemic changes and immune dysregulation are often attributed to increased adipose tissue deposits (p<0.0001, Fig. 5f) and circulating adipokines (Fig. 5g-j)^38,40^. We measured leptin, adiponectin and insulin levels at arrival, diet switch, plus 1-, 2-, and 3-months post-diet switch to determine if perturbations in their typical levels were associated with our phenotypes. After 16 weeks on respective diets, mice fed HFD had significantly elevated leptin, decreased adiponectin, and a lowered adiponectin:leptin ratio, an important indicator of overall adipose tissue inflammation, compared to lean mice (Fig. 5g-j). After diet switch, adipose tissue mass decreased in always obese mice (p<0.0001, Fig. 5f), and metabolically within one month formerly obese mice displayed reduced leptin and insulin while it took two months for adiponectin and three months for the adiponectin:leptin ratio to trend towards the reference point^41^. In summary, when comparing always lean or formerly obese mice to always obese mice, leptin (*p<*0.0001, *p<*0.0001; Fig. 5g), adiponectin (*p<*0.0001, *p=*0.0399; Fig. 5h), and the adiponectin:leptin ratio (*p<*0.0001, *p*=0.0092; Fig. 5i) showed significantly different kinetics, respectively. Insulin (*p<*0.0001, *p<*0.0001; Fig. 5j) levels also varied, suggesting energy maintenance, and potentially metabolic crosstalk to the immune system^42^, was stressed due to the metabolically challenging HFD^43^. This is consistent with previous reports in both preclinical and human data^41,44–46^.

**Figure 5.**
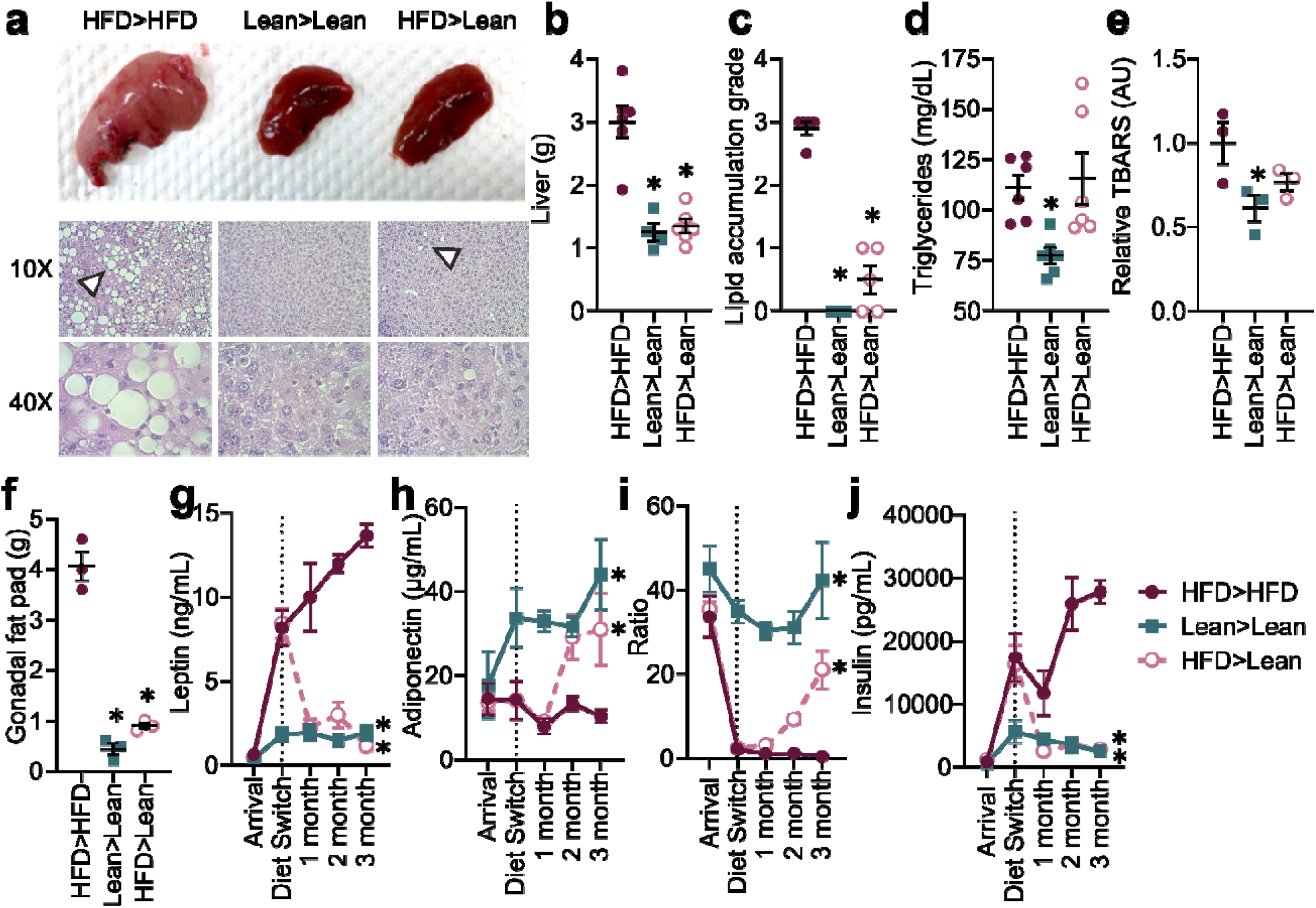
Obesity-related metabolic biomarkers flux due to diet status. (a-f) Systemic effects of high-fat diet on liver (a) representative gross anatomy and fat droplet infiltration visualized by hematoxylin and eosin staining with white arrows signifying fat droplets, (b) mass, (c) overall lipid accumulation at week 30 on diet (18 weeks primary followed by 12 weeks secondary diet), (d) triglyceride content, and (f) oxidative stress. Analysis in (a-f) contains 3-5 mice/group. (g) Mass of gonadal fat pad at week 30 on diet. (g-j) Circulating levels of (g) leptin, (h) adiponectin, (i) adiponectin:leptin ratio and (j) insulin in n=5 mice/group were assayed at arrival (week 0), diet switch (week 18), plus 1-, 2-, and 3-months post-diet switch. Data are represented as means ± standard error and analyzed in (b-f) with ordinary one-way ANOVA with Dunnett’s multiple comparisons test and in (g-j) via two-way ANOVA with Tukey’s multiple comparisons test. Significance represents significant differences compared to always obese (HFD>HFD) mice. Red closed circles = always obese mice, green closed squares = always lean mice, and pink open circles = formerly obese mice.

Given the observed metabolic alterations in our model at time of vaccination, we reasoned that these systemic changes could impair the primary cellular response to immunization (absolute numbers Fig. 6, frequency counts Supplemental Fig. 7). Indeed, we found that mice vaccinated while obese have significantly reduced overall CD4^+^ (p=0.0021, Fig. 6a) and CD8^+^ (p=0.0041, Fig. 6b) T-cell numbers in the spleen while also trending towards reduced B220^+^ B-cell numbers as compared to vaccinated, always lean mice (Fig. 6c). Always lean mice also exhibit increased numbers of CD4^+^ T_EFM_ and CD4^+^ T_CM_ (p=0.0007) cells compared to always obese mice suggesting body weight at vaccination can influence the generation of a CD4^+^ T-cell memory response (Fig. 4). This trend was consistent, albeit not statistically significant (p=0.0831) with an increased frequency in CD4^+^ T_CM_ cells in mice vaccinated while lean as compared to mice vaccinated while obese (Fig. 6d-g). No significant differences were observed between CD8^+^ T_EFM_ and CD8^+^ T_CM_ numbers (Fig. 6e, g). These data highlight how metabolic dysregulation, namely elevated leptin and insulin levels and low adiponectin and adiponectin: leptin levels, at time of influenza vaccination may result in the poor elicitation of a CD4^+^ memory T-cell response, which may explain the poor seroconversion, heightened immunopathology, and reduced survival observed in our formerly obese mice.

**Figure 6.**
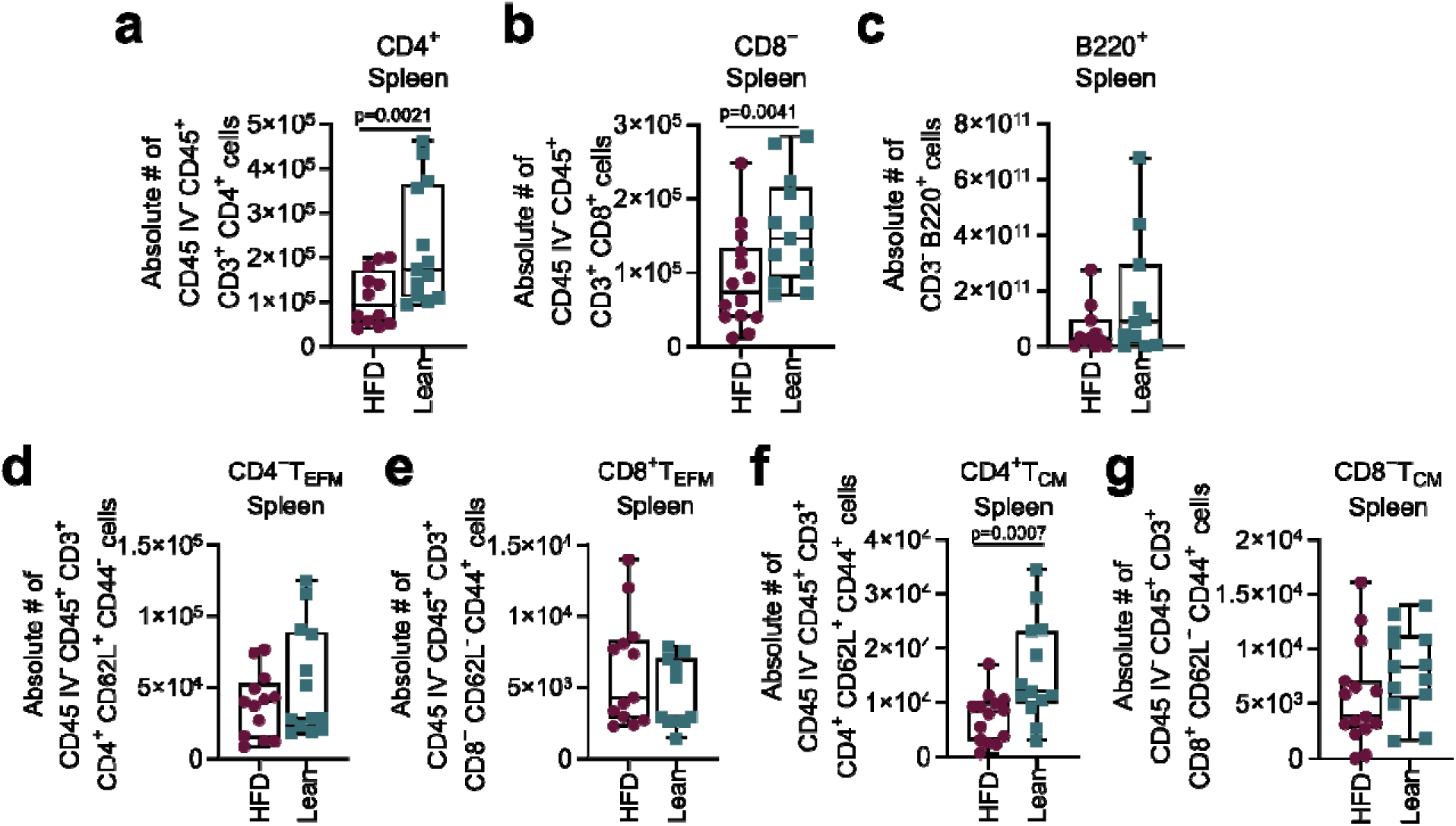
Diet at vaccination defines generation of memory response. Spleens were harvested and homogenized into single-cell suspensions and 1 × 10^6^ cells were stained for CD4^+^ T cells, CD8^+^ T cells, and B220^+^ B cells at 21 days post-vaccination and quantified via flow-cytometry. (a) Absolute number of CD4^+^ and (b) CD8^+^ T cells, (c) B220^+^ B cells, (d) CD4^+^ effector memory (T_EFM_) and (e) CD8^+^ T_EFM_ cells, (f) CD4^+^ central memory (T_CM_) and (g) CD8+ T_CM_ cells. Data are represented as means ± standard error and analyzed using FlowJo version 10.8.1. Outliers in the dataset were removed using the robust regression and outlier removal (ROUT) method (Q=1%). Paired t-test with a two-tailed P-value was used to compare groups at this time point. Note that data include always obese/future formerly obese group (HFD) and always lean (Lean) mice (n = 16 per group) as samples were collected prior to diet switch. Red closed circles = obese and formerly obese mice and green closed squares = lean mice.

### Weight loss occurring pre-vaccination improves viral challenge outcomes and immunological correlates of protection

Mice on HFD for 8 weeks are phenotypically obese but metabolically healthier than mice on our typical 16-week model. They are also better protected from viral challenge post-vaccination despite blunted seroconversion (Supplemental Fig. 8). Thus, we hypothesized that improved metabolic health, even with a history of diet-induced obesity, would be sufficient to preserve vaccine efficacy. To test, formerly obese mice were transferred to control diet for a 4-week wash-out period (Fig. 7a). At this time, weight (p<0.0001, Fig. 7b, c) and metabolic measures, including leptin (p<0.0001, Fig. 7d), are near or approaching the lean baseline. After vaccination, we allowed 3 weeks to elicit vaccine responses on secondary diets and mice were ultimately challenged as before (Fig. 7a). Contrary to weight loss after vaccination (Fig. 1-5), vaccination 1-month post-diet switch significantly improved morbidity in formerly obese mice (Fig. 7e, f; *p<*0.0001, *p<*0.0001) compared to mock vaccinated mice (*p=*0.0715, *p<*0.0001), which translated to mortality (Fig. 7g). Consistent with previous studies, we observed no survival in vaccinated always obese mice and complete survival in vaccinated always lean mice (Fig. 7g, *p=*0.0249), which accompanied decreased viral load at day 7 p.i. In vaccinated always lean and formerly obese mice (Fig. 7h) and increased HAI titers (Fig. 7i). The frequency and numbers of CD4^+^ and CD8^+^ lung T-cells also increased within lung tissue at day 7 p.i. (Fig. 8a, Supplemental Fig. 9a) in formerly obese mice vaccinated after weight loss, which was paired with a reduction in circulating CD4^+^ and CD8^+^ T-cell numbers (Fig. 8b, Supplemental Fig. 9b) as they establish residency within the lung parenchyma. Overall, weight loss before vaccination improved vaccine immunogenicity and efficacy in obese mice. These data highlight the interplay between metabolic health and a well-rounded humoral response and further accentuates the importance of metabolic status at time of vaccination.

**Figure 7.**
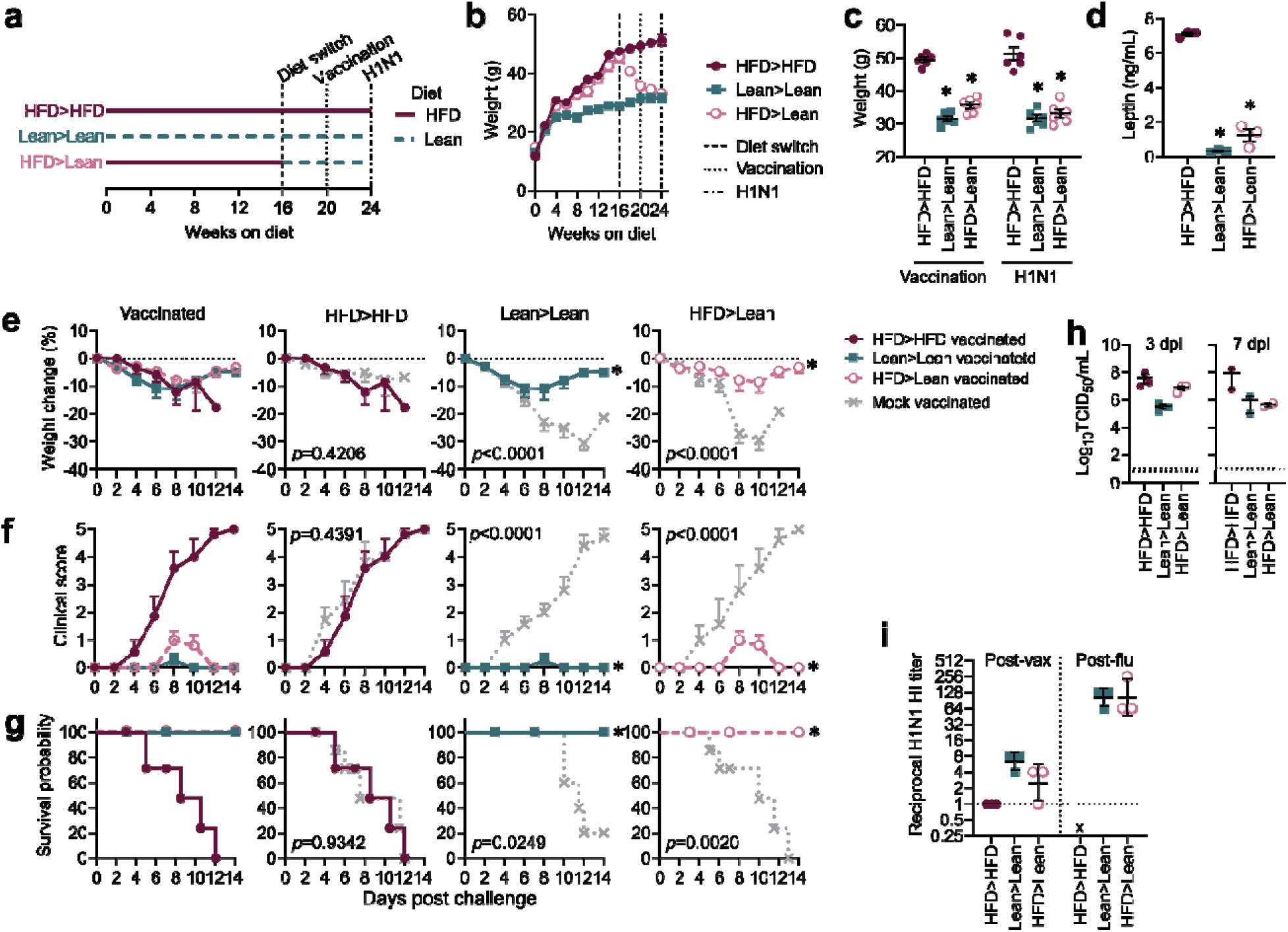
Weight loss prior to vaccination improves survival outcomes. (a) Timeline of vaccination, diet switch, and challenge. (b) Weights at vaccination and challenge with statistical comparisons made using two-way ANOVA with Šídák’s multiple comparisons test between HFD>HFD and indicated groups (n=10 mice/group) at (c) vaccination and challenge. (d) Leptin levels at the time of vaccination in n=3 mice/group. (e-g) Morbidity and mortality in vaccinated mice compared to mock vaccinated mice (n=10 mice/group for HFD>HFD, HFD>Lean, and n=9 Lean>Lean as 1 mouse is excluded from the group due to loss unrelated to study). (e) Weight curves post-challenge with statistical comparisons made via mixed-effects model. (f) Clinical scores post-challenge with statistical comparisons made using a two-way ANOVA. (g) Surviving proportions with censored or event animals indicated by their respective symbols post-challenge and statistical comparisons made using Mantel-Cox log-rank analysis. (h) Lung viral load at day 3 post-challenge compared to always obese, HFD>HFD mice. (i) Hemagglutination inhibition titers at indicated timepoints. Data are representative of three independent experiments and are graphed in (b-f, h) as means ± standard error and in (i) as geometric means ± standard deviation with comparisons made between diet groups to levels in always obese mice (HFD>HFD) via ordinary one- way ANOVA with Dunnett’s correction in (c, g-h). Red closed circles = always obese mice, green closed squares = always lean mice, pink open circles = formerly obese mice, and grey dashed x = mock vaccinated mice.

**Figure 8.**
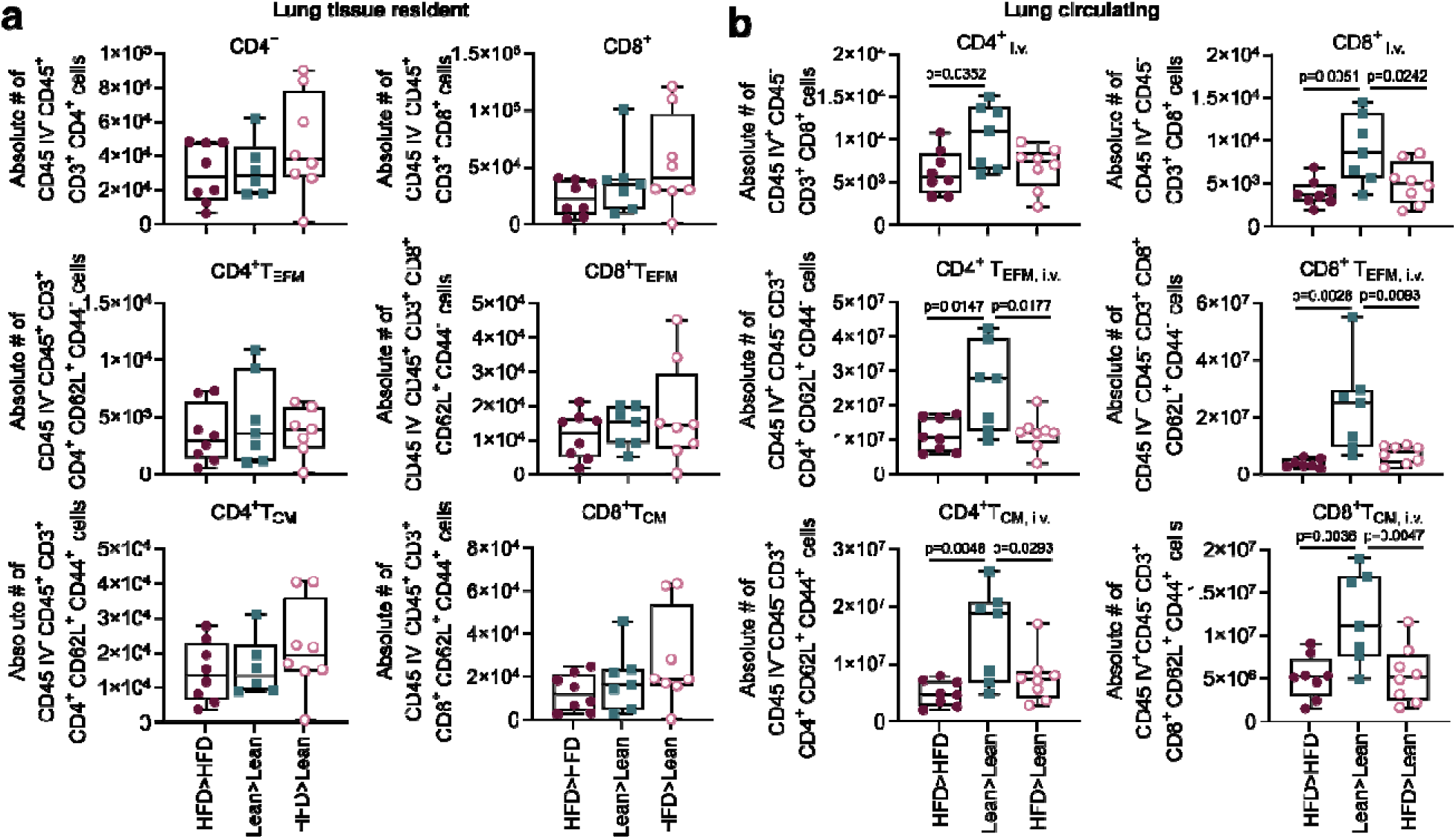
Improved elicitation of memory T cell responses upon weight loss pre- vaccination. Lungs were harvested and homogenized into single-cell suspensions and 1 × 10^6^ cells were stained for CD4^+^ and CD8^+^ T cells at 7 days post-challenge and quantified via flow-cytometry. (a) Absolute number of lung resident CD45_i.v.−_, CD4^+^ and CD45_i.v.−_, CD8^+^ T cells; CD45_i.v.−_, CD4^+^ effector memory (T_EFM_) and CD45_i.v.−_, CD8^+^ T_EFM_ cells; CD45_i.v.−_, CD4^+^ central memory (T_CM_) and CD45_i.v.−_, CD8+ T_CM_ cells. (b) Absolute number of circulating CD45_i.v.−_, CD4^+^ and CD45_i.v.−_, CD8^+^ T cells; CD45_i.v.−_, CD4^+^ effector memory (T_EFM_) and CD45_i.v.−_, CD8^+^ T_EFM_ cells; CD45_i.v.−_, CD4^+^ central memory (T_CM_) and CD45_i.v.−_, CD8+ T_CM_ cells. Data were analyzed using FlowJo version 10.8.1. Outliers in the dataset were removed using the robust regression and outlier removal (ROUT) method (Q=1%). Data include always obese (HFD>HFD), always lean (Lean>Lean), and formerly obese (HFD>Lean) mice (n = 8, n=7, n=7 for each group, respectively) and graphed in as means ± standard error with comparisons made between diet groups via two-way ANOVA. Red closed circles = always obese mice, green closed squares = always lean mice, and pink open circles = formerly obese mice

## Discussion

Our current, standard-of-care influenza vaccinations are poorly immunogenic in high-risk hosts, including the aged, pregnant, and those with metabolic dysfunction^47^. Preclinical studies suggest obese mouse models present with a reduced breadth and depth of the antibody response^16^, reduced generation of innate, effector and memory facets of cellular immunity^13,17,48^, and poor responsiveness to adjuvants upon vaccination^16^, while epidemiological evidence points towards poor immunigenicity^15,49^ and increased waning of vaccine-induced protection^15^ in obese populations compared to lean. While novel vaccine platforms and adjuvants may provide increased protection to this and other high-risk groups, a recurrent open question is the feasibility of host-directed measures to improve immunization efficacy. Here, we show weight loss failed to improve survival of formerly obese mice at viral challenge if occurring post-vaccination (Fig. 1, Supplemental Fig. 1), yet weight loss pre-vaccination resulted in 100% of formerly obese mice surviving challenge (Fig. 7). Essentially, the snapshot of diet group and associated metabolic dysfunction captured at the time of vaccination fixed the recall response to viral challenge, even after diet change and subsequent weight loss.

Appropriately timed innate responses permit a cascade of adaptive and repair responses to follow, thus limiting severe sequelae upon infection. While blunted immune responses are characteristic of obese hosts, paradoxically this delay may lead to immunopathology at later timepoints^5051^. Previous studies have interrogated how weight loss may change these dynamics^52, 3417^. In line with our study, Rebeles et al. show weight loss in DIO mice fails to improve survival or effector T-cell numbers and recall upon secondary influenza challenge^34^. Our study extends this work and demonstrates that at time of vaccination, exposure to HFD can impair the generation of a primary memory CD4^+^ T-cell response, as evidenced by decreased numbers of overall CD4^+^ T-, CD4^+^ T_EFM,_ and CD4^+^ T_CM_ cells in the spleens of our diet group exposed to an obesogenic protocol. These findings are in alignment with the blunted seroconversion we observed in mice vaccinated when on HFD, highlighting how nutritional status at the time of immunization influences primary memory T-cell generation and shapes broader humoral immunity at the time of recall.

While formerly obese mice did show reduced viral spread and modestly improved interferon responses due to their “corrected” nutritional status at challenge, we posit their primary response was “fixed” due to immunization occurring while under HFD conditions. Consistent with this hypothesis, formerly obese mice show an improvement in memory T-cell numbers at 7 dpi within the lung and in circulation, but it is not sufficient to overcome viral-induced damage. Given the limitations posed by the lack of survivors in diet groups exposed to HFD, we focused on investigating T-cell responses solely at one time point that would ideally capture a cellular recall response to challenge (day 7 p.i.)^35^. While this hinders our ability to evaluate infiltrating T-cell kinetics in each of our diet groups within our study, it warrants future studies that define the dynamics of T-cell infiltration possibly using a lower infection dose to circumvent survivor bias. Additionally, functional studies evaluating virus-specific T-cell interactions would be beneficial to illustrate whether homing of T-cells into the bona fide tissue effector site, the lung, during influenza infection is reduced in groups exposed to HFD at time of immunization, which could explain why an increase of T-cells post-challenge in mice vaccinated while obese is still not sufficient to combat viral-induced lysis of infected cells as opposed to mice vaccinated when lean. For always lean mice, the training of the immune system was done while fed a standard chow, thus supporting the generation of quality effector memory T-cell response (Fig. 6) which reduced immunopathology (Fig. 2) and improved effectiveness of a cellular recall to the lung at challenge (Fig. 3, 4), all culminating in high survival probability (Fig. 1, Supplemental Fig. 1).

Adipokines, especially leptin, are implicated in attenuating both innate and adaptive immune responses in high-risk populations and are highly responsive to diet status^53,54^. Our HFD was 60% fat, with lard as the primary source. Lard is high in saturated fatty acids, well-documented stimulators of toll-like receptor-dependent inflammatory processes^55–57^. The diets contained 20% carbohydrates with nearly a third of total carbohydrates being sucrose–much higher than the 3% sucrose in the lean chow^58,59^. Sucrose is a disaccharide of glucose and fructose, which both contribute to metabolic abnormalities when consumed in excess^60–63^. Excess sugars can be converted to fatty acids, which are stored in the liver and overtime result in reactive oxygen species formation, systemic meta-inflammation, and contribute to imbalanced adipokine production (Fig. 5)^62–64^. The pro-inflammatory milieu stemming from exposure to HFD may have blunted both T-cell memory formation at the time of vaccination as well as the innate and adaptive response to H1N1 virus at challenge^65^. Previous studies of weight loss and vaccination have demonstrated the intractable nature of adipokine dysfunction in the obesogenic state, especially during influenza infection^34,66,67^. In murine models, even with weight loss and return to a metabolically healthy baseline, HFD generates persistent macrophage and CD8^+^ T-cell-mediated inflammation in the liver, adipose tissue, and lung as well as impaired T-cell responsiveness to infection^20,34,68^. These prolonged effects of obesity have been noted in rats, macaques, and humans, and in tissues as varied as skeletal muscle^69^ and tumors^70^ and in offpsring^71,72^, indicating diet’s effects on immunometabolism may persist long after weight loss.

While our study successfully establishes a link between diet at time of vaccination and survival at challenge, we had several limitations. First, we were unable to determine the kinetics of the primary cellular response post-vaccination and the memory recall response post-infection as our sample sizes limited our analysis to a snapshot of the immunological infiltrates at a single timepoint. Similarly, we are unable to determine the timepoint at which mice exposed to HFD reach a “point of no return” metabolically. Our work suggests 12-weeks is sufficient to yield the phenotypic, metabolic, and immunologic consequences of HFD and related metabolic syndrome on vaccination; however, 8-weeks of HFD does not reduce influenza vaccine efficacy. We did not explore finer investigation into the kinetics of adipose tissue dysfunction that arises post-HFD exposure, including immunological measures within the adipose tissues as well as systemic metabolic dysfunction stemming from the endocrine functions of adipocytes^73,74^. Previous studies have displayed varying effects of HFD and the length of diet exposure on the adipose tissue secretome^75,76^, adipose tissue resident immune cells^77,78^, and various classes of lymphocytes in general^17,79^. In our studies, diet status at the time of vaccination correlated with cellular recall responses at challenge, the resulting lung inflammation, and overall survival trends while diet group at challenge was related to viral spread and interferon responses. It is known nutrition and dietary metabolites can influence the cellular metabolism of immune cells^34,80^ and our results necessitate study into the role bioactive dietary components play on the innate and cellular immune responses at both immunization and infection^55,81,82^.

The relationship between obesity and infectious diseases is not unique to influenza virus. Epidemiological data and studies in obese mice demonstrate obesity is a risk factor for severe COVID-19^26,83^ and other infectious diseases^84,85^, and weight loss has been associated with improved outcomes^86^. Additionally, other host-directed interventions may improve vaccine efficacy. Clinical trials suggest exercise can boost antibody and cellular-mediated responses upon vaccination, especially in high-risk groups such as the elderly and the obese^87–89^. Future studies must be performed to untangle how components of the obesogenic diet, resulting lung microenvironment, and other host behaviors impact formation of primary and recall responses upon exposure to the pathogen-of-interest, as well as the efficacy of various vaccine platforms, novel metabolically-targeted adjuvants, and timing of vaccine boosters in these high-risk hosts^90^. Understanding how nutritional status influences immunization responses is important not only for those under- or over-nourished, but also increasingly of concern with weight cycling and extreme metabolic flux, fad diets limiting both macro and micronutrients, and climate change or other global logistical concerns threatening the local availability of food.

## Acknowledgements

The authors gratefully acknowledge the expert technical assistance and scientific discussions from Dr. Ericka Roubidoux, Kristin Wiggins, Brandi Clark, members of the Schultz-Cherry and Thomas labs, and pathological expertise of Dr. Peter Vogel. We also thank Derek Warren, Crystal Oudomvilay, and Chava Roberts for assisting in sample and assay preparation. Thank you to the St. Jude Animal Resource Center and Veterinary Pathology Core for their helpful experimental assistance. This work was supported by ALSAC, the National Institute of Allergy and Infectious Diseases (NIAID) at the National Institutes of Health (NIH) under Department of Health and Human Services (HHS) contract HHSN27220140006C for the St. Jude Center of Excellence for Influenza Research and Surveillance (CEIRS), by the NIAID Collaborative Influenza Vaccine Innovation Centers (CIVIC) contract 75N93019C00052, contract 75N93021C00016 for the St. Jude Center of Excellence for Influenza Research and Response (CEIRR) to S. S.-C. and P. G. T. and NIAID R01 AI140766-03 to S.S.-C. A.V.P. was supported by the St Jude Graduate School for Biomedical Sciences and NIAID NIH Award Number F31AI161986. The content is solely the responsibility of the authors and does not necessarily represent the official views of the National Institutes of Health.

## Author contributions

Conceptualization - R.H., A.V.P., S.S-C.; Methodology - R.H., A.V.P., S.S-C. P.G.T.; Validation - R.H., A.V.P., S.S-C.; Formal analysis - R.H., A.V.P., R.C.S.; Investigation - R.H., A.V.P., B.L., V.H., B.S., S.C., L.A.V.D.V., R.C.S.; Resources - R.H., A.V.P., B.L., B.S.,L.A.V.D.V., P.G.T., S.S-C.; Data curation - R.H., A.V.P.; Writing - original draft - R.H., A.V.P.; Writing - review and editing - All authors; Visualization - R.H., A.V.P.; Supervision - S.S-C.; Project administration - R.H., A.V.P, S.S-C.; Funding acquisition - S.S-C. and P.G.T. R.H. and A.V.P. share co-first authorship and contributed equally to this work.

## Materials & Correspondence

Any requests for reagents, materials, or correspondence should be directed to the corresponding author at.

## Data Availability

All data generated or analyzed during this study are included in the published article and its supplementary information files.

## Methods (1594)

### Animal husbandry and ethics

All procedures (protocol 513) were approved by the St. Jude Children’s Research Hospital Institutional Animal Care and Use Committee (IACUC) and complied with the Guide for the Care and Use of Laboratory Animals. These guidelines were established by the Institute of Laboratory Animal Resources, approved by the Governing Board of the U.S. National Research Council, and has an approved Animal Welfare Assurance Statement on file with the Office of Laboratory Animal Welfare (A3077-01). Mice were maintained at 12 h light-dark cycles at ambient temperature 68°F and 45% humidity.

Designated diet and water were provided *ad libitum* for the duration of the experiments. When necessary due humane endpoints or timepoint collection, mice were humanely euthanized following AVMA guidelines.

### Diet and vaccination schedules

Male and female 3-week-old C457Bl/6 mice (JAX) were randomized to control diet (Lab Diets, #5001, 15% fat, 30% protein, 55% carbohydrate) or high-fat diet (HFD, Research Diets #D12492, 60% fat, 20% protein, 20% carbohydrate). Weights were measured biweekly, and serum collected at arrival and pre-vaccination. After 2 or 4 months on the respective diets, mice were immunized with 50 μL of β-propiolactone (BPL)-inactivated A/California/04/2009 (H1N1) virus, standardized to 1 μg hemagglutinin (HA) protein per mouse in phosphate-buffered saline (PBS), administered in the right quadriceps femoris muscle. Mock-vaccinated mice received 50 μL of PBS. We allowed 2 weeks to generate a vaccination response, then mice were randomized to receive either the opposing diet or maintained on their primary diet for the duration of the study. Monitoring of weights continued biweekly and serum collections monthly for 1 or 3 months (Fig. 1a, Supplemental Table 1). For studies concerning weight change prior to vaccination, mice were randomized to control diet or HFD as before. After 4 months, half were switched to the opposing diet for 1 month prior to vaccination and were ultimately challenged 1 month after vaccination.

### Viral titer determination

Viral titer was determined as previously reported^91^. Briefly, confluent Madin-Darby canine kidney (MDCK) cells were inoculated in triplicate with serial 10-fold dilutions of whole lung homogenates (bead beat in 500 μL PBS) in 100 μL of MEM plus 0.75% BSA and TPCK-trypsin. After 3 days of incubation at 37°C, 50 μL of supernatant was mixed with 50 μL of 0.5% packed turkey red blood cells diluted in PBS for 45 minutes at room temperature. Observance of hemagglutination indicated a positive result. Quantitative tissue culture infectious dose-50 (TCID_50_) determined via the Reed–Muench method. A mean lethal dose-50 (MLD_50_) was previously calculated for the viral stock in naive, wild-type C57BL/6 male and female mice to standardize inoculums.

### Viral challenge

At indicated time points, mice were challenged at a 10X MLD_50_ dose (determined in preliminary studies) with A/California/04/2009 (H1N1) virus in 25 μL PBS administered intranasally under light anesthesia (3-4% inhaled isoflurane). Weights and clinical scores were recorded for 14 days. Moribund mice that had lost more than 30% body weight and/or reached clinical scores of greater than 3 were humanely euthanized.

Clinical signs were scored as follows: 0=no observable signs, 1=active, squinting, hunching or scruffy appearance; 2=squinting and hunching, active only when stimulated; 3=excessive hunching, squinting, not active when stimulated; 4=hind-limb paralysis, shivering, rapid breathing, moribund and 5=death. Clinical signs were agreed upon by at least two experienced researchers. At days indicated, mice were euthanized and whole lung, spleen, serum, or other indicated tissues harvested and processed immediately or stored at -80°C for future analysis. All results are representative of 2 independent experiments.

### Metabolic biomarker ELISA

Whole blood was collected, allowed to clot at room temperature, then centrifuged at 5,000 x g for 15 min to separate sera and immediately snap-frozen and stored at -80°C. MesoScale Discovery metabolic assay platform enzyme-linked immunosorbent assay (ELISA) was used to quantify protein concentrations of c-peptide, total ghrelin, total GLP-1, insulin, leptin, and total protein tyrosine tyrosine in sera (MSD; cat#K15301K).

Sera were diluted 1:2 in PBS and used per manufacturer’s protocol. The adiponectin/acrp30 DuoSet ELISA (1:10,000 dilution, R&D Systems, cat#DY1119) and leptin DuoSet ELISA (1:5 dilution, R&D Systems, cat#DY406) were used to measure and confirm adiponectin and leptin levels, respectively. Corrected OD values were compared to the standard curve to calculate total concentration. Corrected OD values were compared to the standard curve to calculate total concentration. To estimate oxidative stress, total thiobarbituric acid reactive substances (TBARS) in liver specimens was assessed with colorimetric TBARS assay kit (Cayman Chemical, #10009055) according to the manufacturer’s suggestions.

### RNA extraction and RT-qPCR

Bead beat whole lung homogenate (100 μL) was further dissociated in 1 mL of TRIzol reagent then centrifuged at 500 x g for 3 minutes to pellet cellular debris. Supernatant underwent phenol-chloroform RNA extraction per manufacturer’s protocol (Invitrogen, cat# 15596026). Eluted RNA was resuspended in 50 μL of nuclease-free water and quantified spectrophotometrically (Nanodrop). RNA was converted to cDNA using the SuperScript IV VILO kit. A 1:20 dilution of cDNA was then used in subsequent quantitative PCR on using the QuantiFast platform and QuantiTect primer assays for *Ifna2* (NM_010503), *Ifnb1* (cNM_010510), *Ifnl2* (NM_001024673), and *Gapdh* (NM_008084). Gene expression was standardized to endogenous *Gapdh* cycle thresholds (ΔCt) and ΔΔCt values calculated in relation to HFD>HFD always obese mice.

### Histopathology

Deeply anesthetized mice were perfused with 10% neutral buffered formalin and tissues collected at the indicated time post-infection. Preserved lung tissues were embedded in paraffin, sectioned, and stained with a customized *in situ* hybridization (ISH) probe specific for the A/California/04/2009 (H1N1) virus NS1 gene (GenBank accession #JF915191; Advanced Cell Diagnostics), to identify active viral replication. Lung slides were also stained with hematoxylin and eosin (H&E) for pathological analysis with total pathological scores compared across diet groups and days post-infection through z-score standardization by pathology. Livers were collected in a similar manner and stained with H&E to visualize lipid accumulation with images captured on an Axio Lab A.1 microscope fitted with an AxioCam ERc5s (Zeiss) using the ZEN 3.0 software. The extent of lipid infiltration is the average score of two blinded, independent researchers (R.C.S. and R.H.)^92^.

### Antibody quantification

To quantify influenza-specific antibodies, mouse sera were treated with receptor-destroying enzyme (RDE; Seiken; cat#370013) and hemagglutination inhibition (HAI) and microneutralization (MN) assays were performed as described^16^. Total IgG levels were assayed according to manufacturer’s instructions (Thermofisher; Cat # 88-50400- 22). Viral-specific IgG^93^ were quantified by coating a 96-well plate with whole A/California/04/2009 (H1N1) virus overnight at 4°C, bocking in 5% nonfat dry milk, then adding indicated dilutions of RDE-inactivated sera in duplicate for 1 hour at room temperature. After washing, horseradish peroxidase (HRP) conjugated anti-mouse IgG was added for 1 hour at room temperature, after which detection and quantification was enabled by addition of 3,3’,5,5’-tetramethylbenzidine (TMB). Data are represented as arbitrary units as the difference in absorbance (450 nm-570 nm).

### Flow cytometry for lung and spleen memory T-cells

Mice were lightly anesthetized with 3% inhaled isoflurane, then retro-orbitally injected with 2 μg/mouse of APC anti-mouse CD45 (Biolegend, cat#103112; clone 30-F11) antibody in 50 μL PBS for intravenous labeling of CD45^+^ circulating lymphocytes 15 minutes prior to euthanization. Mouse lungs were excised and finely minced with a surgical blade, then digested in 1X HBSS medium containing 1 mg/mL DNase I (Worthington Biochem.com, cat#LS002145) and 2.5 mg/mL Collagenase Type I (Worthington Biochem.com, cat#LS004196) at 37°C for 30 min. The digest was dissociated with a 1mL pipette tip and passed through a 100 μm filter into a 50 mL tube.

Any additional pieces of the lung were dissociated using a 10mL syringe plunger and washed using 20 mL of DMEM/F12 (Corning; cat#10-092-CV). Cells were centrifuged at 300 x g for 5 min at 4°C and treated with 1 mL of Ammonium-Chloride-Potassium (ACK) Lysing Buffer (Gibco; cat#A1049201). After 2 minutes, 10 mL DMEM/F12 was added, and cells pelleted at 300 x g for 5 min at 4°C. Cells were resuspended in 1 mL DMEM/F12 and automated cell counts recorded (Cellometer). For antibody staining, 1×10^6^ cells were plated and then incubated with Fc block (Biolegend; cat#101320; 1:100 dilution) and surface stained for 20 min at 4°C in the dark. Cells were washed one time with FACS buffer and then resuspended in 100 μL of an antibody cocktail containing the following: murine anti-CD45_i.v.+_ (Biolegend, cat#103112; clone 30-F11; 1:100 dilution), anti-CD45_i.v.-_ (Biolegend; cat# 103154; clone 30-F11; 1:200), anti-CD4 (TONBO Biosciences; clone GK1.5; 1:200 dilution), anti-CD8 (TONBO Biosciences; cat# 35-0081-U500; clone 53-6.7; 1:200 dilution), anti-CD44 (TONBO Biosciences; cat#75-0441-U100; clone 1M7; 1:100 dilution), anti-CD3 (Biolegend; cat#100246; clone 17A2; 1:100 dilution), anti-CD62L (Biolegend; cat#104428;clone MEL-14;1:100 dilution), anti-CD69 (Invitrogen; cat#45-0691-82;clone H1.2F3; 1:100 dilution), anti-CD103 (Biolegend, cat#121426; clone 121426; 1:100 dilution) and a viability dye, Ghost Dye 510 (TONBO Biosciences; cat#13-0870-T100; 1:100 dilution) antibodies. Additionally, cells were also stained with a separate cocktail that included anti-B220 (Biolegend; cat#103240; clone RA3-6B2; 1:100 dilution) and a viability dye, Ghost Dye 510 (TONBO Biosciences; cat#13-0870-T100; 1:100 dilution) antibodies. Cells were incubated with the antibody cocktails for 30 minutes at 4°C in the dark, and then washed twice with FACS buffer before resuspending in a final volume of 100 μL. Cells were analyzed using a BD LSRII Fortessa and BD FACSDIVA software (Becton Dickinson). Data were analyzed using FlowJo software (FlowJo v10.8.1).

### Statistical analysis

Prior to animal studies, power analysis was performed to ensure study rigor with β=0.2^94,95^. Data were compiled and organized using Microsoft Excel, FlowJo v10.8.1, and GraphPad Prism v9.3.1 with figures finalized in Inkscape vector graphics editor. Statistical analysis was performed using GraphPad Prism v9.3.1 as described in the figure legends. When required due to experimental timepoints, survival proportions include data up to timepoints for any censored mice (Supplemental Table 1). Statistical significance was defined as α=0.05 and, unless otherwise specified, is comparing the always obese HFD>HFD diet cohort to other experimental groups.

