## Supplemental Tables and Figures for "Improved metabolic syndrome and timing of weight loss is crucial for influenza vaccine-induced immunity in obese mice"

**Supplementary material**

**Supplemental Table 1. Summary of studies and sample sizes per group.**

| Study | Sex | N | Total groups | Vaccine and Infection | Design^a^ | Outcomes |
| --- | --- | --- | --- | --- | --- | --- |
| 1 | Male | 8^b^-10 | 6 | BPL-inactivated A/CA/09/2009^c^ vaccine or mock vaccination; A/CA/04/2009 challenge | Vaccine at week 8, diet switch at week 10, challenge at week 14 | Sera collected pre- and post-vaccine and challenge, survival and morbidity assessed |
| 2 | Male | 10 | 4 | BPL-inactivated A/CA/09/2009 vaccine; A/CA/04/2009 challenge | Vaccine at week 16, diet switch at week 18, challenge at week 30 | Sera collected pre- and post-vaccine and challenge, survival and morbidity assessed, viral loads and qPCR at 3 dpi^d^, flow cytometry trial at 10 dpi |
| 3 | Male | 10 | 4 | Mock vaccine; A/CA/04/2009 challenge | Mock vaccine at week 16, diet switch at week 18, challenge at week 30 | Sera collected pre- and post-vaccine and challenge, survival and morbidity assessed, viral loads and qPCR at 3 dpi, flow cytometry trial at 10 dpi |
| 4 | Male | 10 | 4 | BPL-inactivated A/CA/09/2009 vaccine; A/CA/04/2009 challenge | Vaccine at week 16, diet switch at week 18, challenge at week 22 | Sera collected pre- and post-vaccine and challenge, survival and morbidity assessed, viral loads, qPCR, and histology at 3 dpi, flow cytometry trial at 10 dpi |
| 5 | Female | 5 | 8 | BPL-inactivated A/CA/09/2009 vaccine or mock vaccine; A/CA/04/2009 challenge | Vaccine at week 16, diet switch at week 18, challenge at week 22 | Sera collected pre- and post-vaccine and challenge, survival and morbidity assessed |
| 6 | Male | 10 | 4 | Mock vaccine; A/CA/04/2009 challenge | Vaccine at week 16, diet switch at week 18, challenge at week 22 | Sera collected pre- and post-vaccine and challenge, survival and morbidity assessed, viral loads, qPCR, and histology at 3 dpi, flow cytometry trial at 10 dpi |
| 7 | Male | 10 | 8 | BPL-inactivated A/CA/09/2009 vaccine or mock vaccination; A/CA/04/2009 challenge | Diet switch at week 16, vaccine (or mock) at week 20, challenge at week 24 | Sera collected pre- and post-vaccine and challenge, survival and morbidity assessed |
| 8 | Male | 10 | 4 | BPL-inactivated A/CA/09/2009 vaccine; A/CA/04/2009 challenge | Vaccine at week 16, diet switch at week 18, challenge at week 22 | Sera collected pre- and post-vaccine and challenge, survival and morbidity assessed, histology at 5 dpi, viral loads at 7 dpi |
| 9 | Male | 10 | 8 | BPL-inactivated A/CA/09/2009 vaccine; A/CA/04/2009 challenge | Vaccine at week 16, diet switch at week 18, challenge at week 22 | Sera collected pre- and post-vaccine and challenge, lungs and spleens collected for flow cytometry at day 21 post vaccination or 7 dpi |
| 10 | Male | 10 | 8 | BPL-inactivated A/CA/09/2009 vaccine; A/CA/04/2009 challenge | Diet switch at week 16, vaccine at week 20, challenge at week 24 | Sera collected pre- and post-vaccine and challenge, lungs and spleens collected for flow cytometry at day 21 post vaccination or 7 dpi |

^a^Weeks are measured from start of original study diet beginning at week 3 of age

^b^Pilot study 1 included 8 mice for HFD>HFD group and 10 mice for all other groups and included only a Lean>Lean and HFD>HFD unvaccinated control

^c^A/California/04/2009 (H1N1) virus was propagated in-lab through passage in MDCK cells and inactivated via addition of β-propiolactone

^d^dpi=days post infection for the viral challenge

**Supplemental figures and legends**

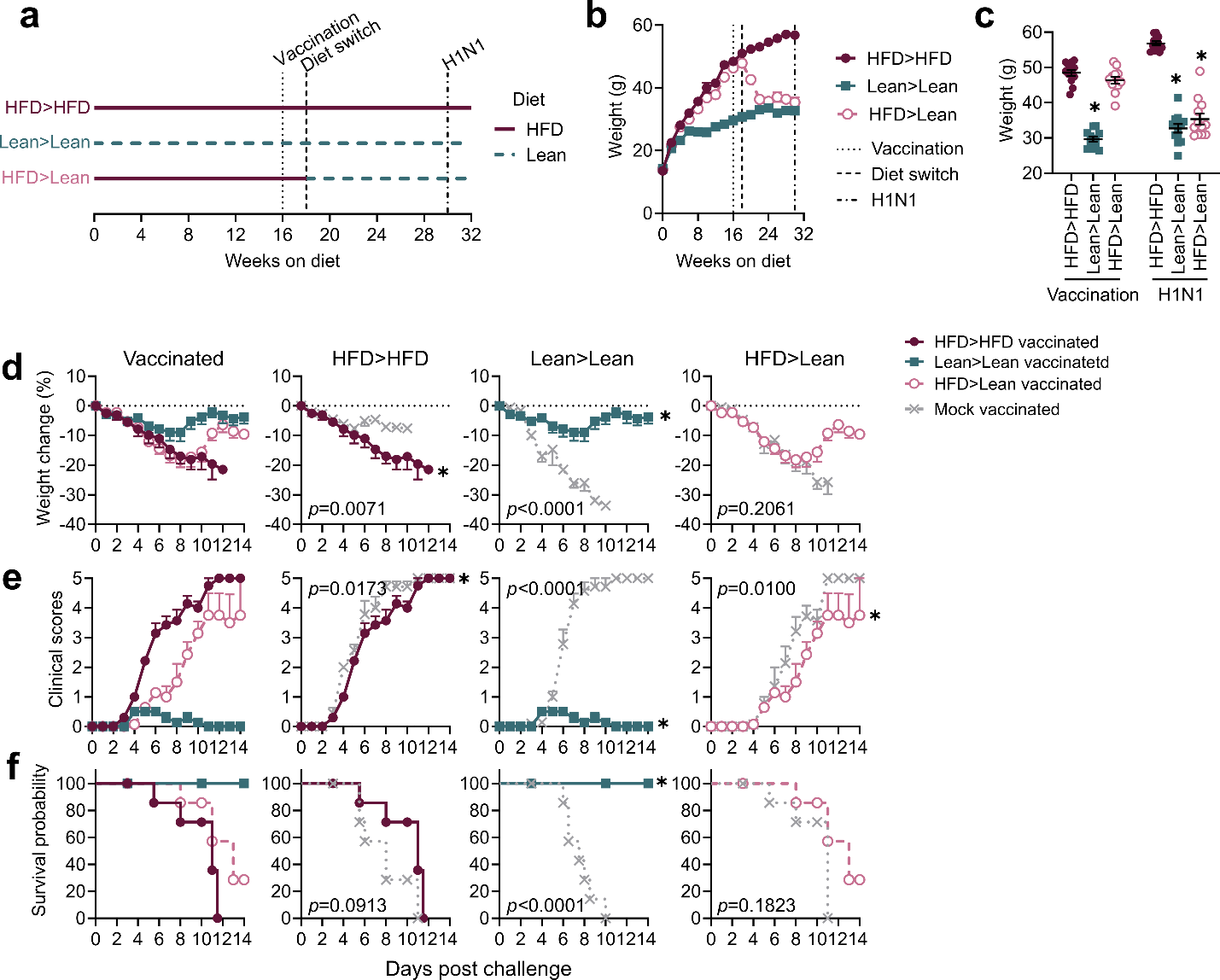

**Supplemental Figure 1. Long-term weight loss does not improve survival upon viral challenge.** (a) Timeline of diet administration, vaccination, and challenge for 12-week long-term diet protocol. (b) Weights of mice through short-term diet protocol and at (b) vaccination and challenge. (d-f) Morbidity and mortality in vaccinated mice compared to mock vaccinated mice with n=10 mice/group. (d) Weight curves post-challenge with statistical comparisons made via mixed-effects model. (e) Clinical scores post-challenge with statistical comparisons made using a two-way ANOVA. (f) Survival post-challenge with statistical comparisons made using Mantel-Cox log-rank analysis. Data in (b-e) are representative of two independent experiments and are graphed as means ± standard error and in (f) as surviving proportions with censored or event animals indicated by their respective symbols. Statistical comparisons test between HFD>HFD and indicated diet group (n=10 mice/group) in (b, c) and between mock and vaccinated animals within a single diet group in (d-f). Red closed circles = always obese mice, green closed squares = always lean mice, pink open circles = formerly obese mice, and grey dashed x = mock vaccinated mice.

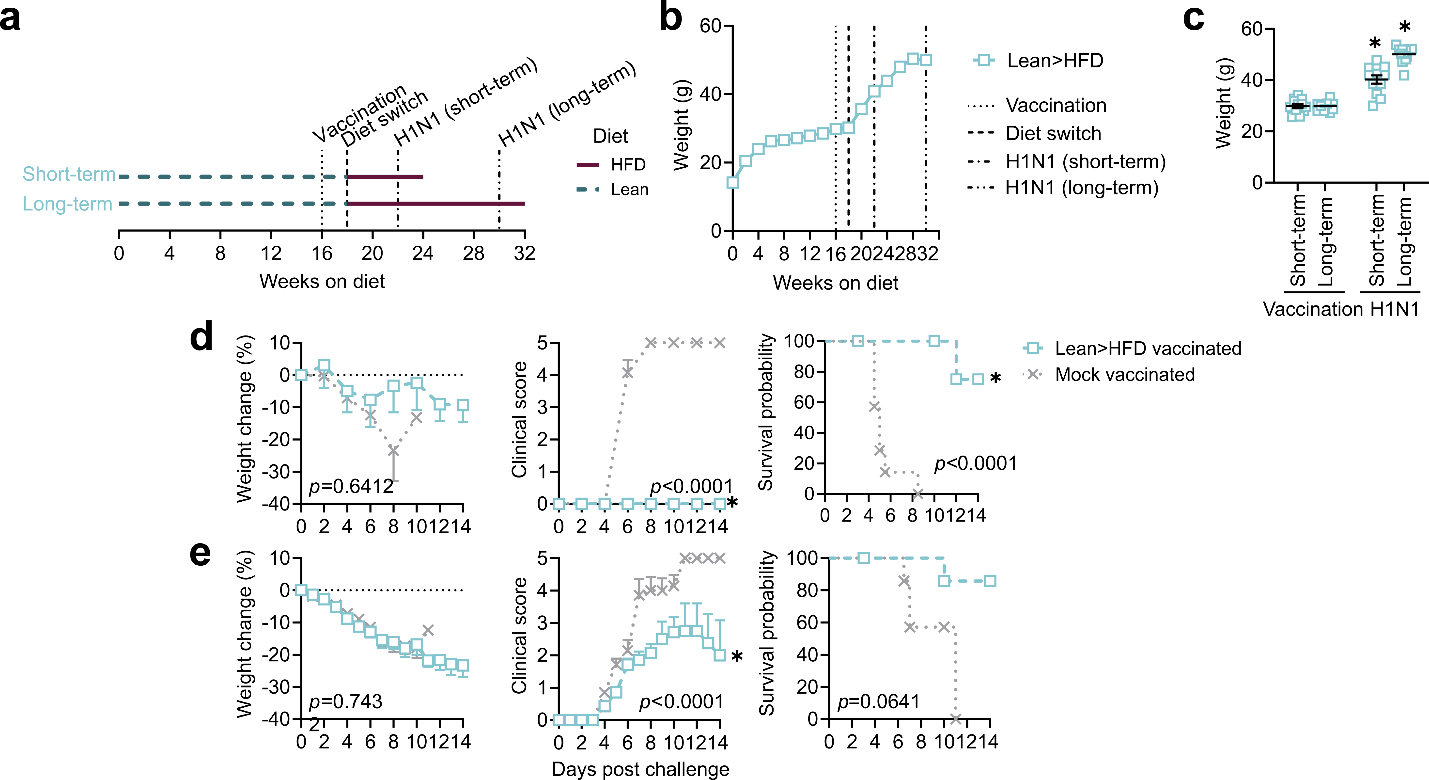

**Supplemental Figure 2. Weight gain post-vaccination has modest impacts on survival at challenge.** (a) Timeline of diet administration, vaccination, and challenge for short- and long-term weight gain protocols. (b) Weights of mice on short- and long-term weight gain protocol at (c) vaccination and challenge. (d-e) Morbidity and mortality in vaccinated mice compared to mock vaccinated mice with n=10 mice/group. (d-e) Weight curves post-challenge with statistical comparisons made via mixed-effects model. Clinical scores post-challenge with statistical comparisons made using a two-way ANOVA. Survival post-challenge with statistical comparisons made using Mantel-Cox log-rank analysis. Data are representative of three independent experiments and are graphed as means ± standard error and, for survival graphs, as surviving proportions with censored or event animals indicated by their respective symbols. Statistical comparisons test between HFD>HFD and indicated diet groups (n=10 mice/group) in (b, c) and between mock and vaccinated animals within a single diet group in (d, e). Light blue open squares = formerly lean mice and grey dashed x = mock vaccinated mice.

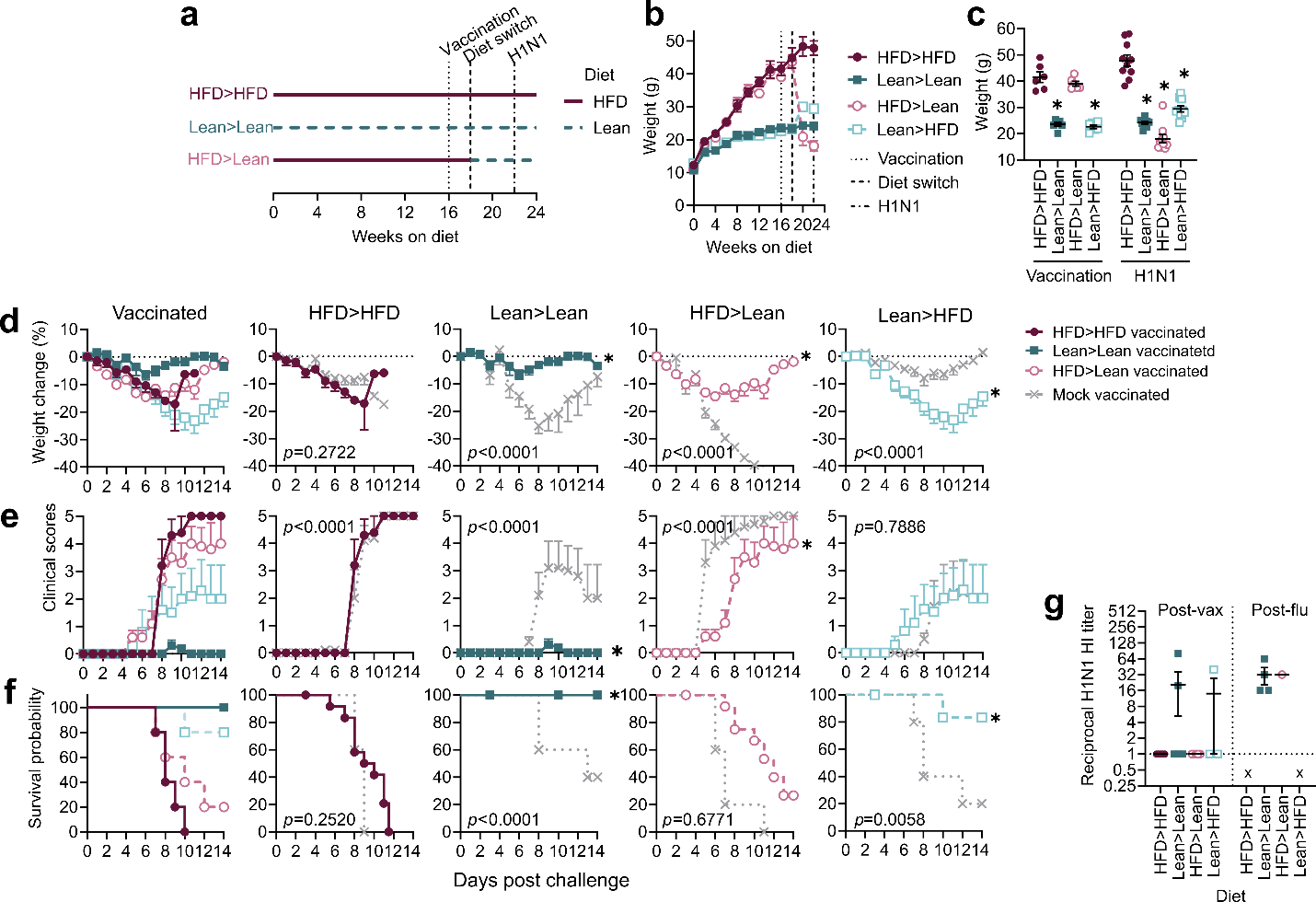

**Supplemental Figure 3. Similar responses to challenge post-diet switch in female mice.** (a) Timeline of diet administration, vaccination, and challenge in short-term diet and weight gain protocols (b) Weights of female mice on diet and weight gain protocols at (c) vaccination and challenge. (d-f) Morbidity and mortality in vaccinated mice compared to mock vaccinated mice. (d) Weight curves post-challenge with statistical comparisons made via mixed-effects model. (e) Clinical scores post-challenge with statistical comparisons made using a two-way ANOVA. (f) Survival post-challenge with statistical comparisons made using Mantel-Cox log-rank analysis. Data in (b-e) are represented as means ± standard error and in (f) as surviving proportions with censored or event animals indicated by their respective symbols. (g) Effector antibody levels in sera collected and quantified by hemagglutination inhibition (HAI). Mice assayed longitudinally with n=5 per group except for the day 21 post-influenza challenge timepoint in which only surviving mice were assayed. Data displayed as geometric means ± standard deviation with x indicating no mice surviving to time point. Statistical comparisons test between HFD>HFD and indicated diet group (n=10 mice/group) in (b, c, g) and between mock and vaccinated animals within a single diet group in (d-f). Red closed circles = always obese mice, green closed squares = always lean mice, pink open circles = formerly obese mice, light blue open squares = formerly lean mice and grey dashed x = mock vaccinated mice.

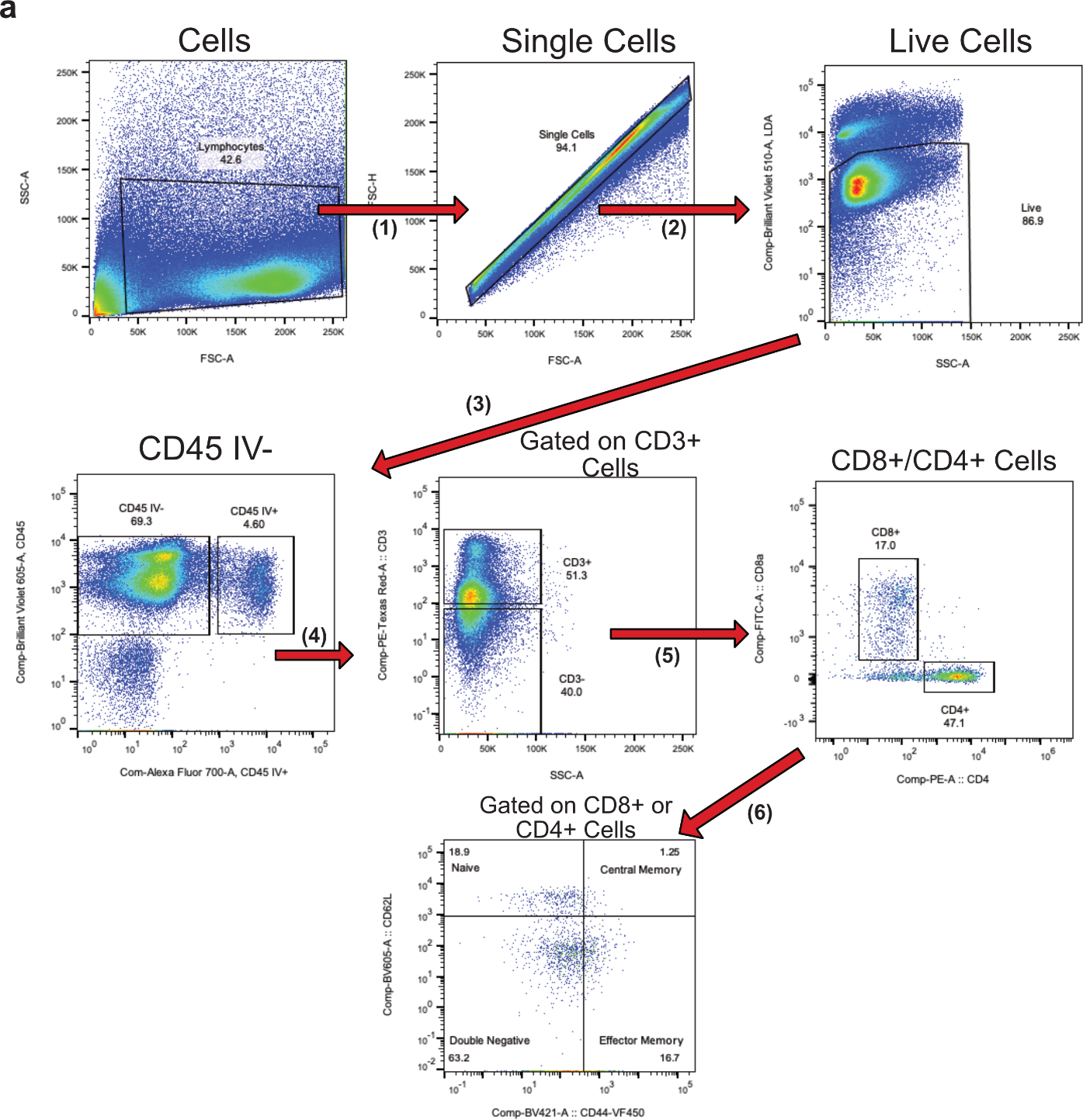

**Supplemental Figure 4. Assessment of T-cell surface phenotype in spleen and lungs upon distinct diet regimens in male mice post-vaccination and post-influenza challenge.** (a) Gating strategy for identifying live CD45^+^CD45 IV^-^ CD3^+^ cells in mouse spleen and lung samples. (b) Each flow plot represents cells gated on live CD45^+^CD45 IV^-^ CD3^+^ from an individual sample as indicated in the gating strategy. Naïve (CD62L^+^CD44^-^), Central Memory (T_CM_, CD62L^+^CD44^+^), and Effector Memory (T_EFM_, CD62L^-^CD44^+^) cells are labeled on representative flow plots.

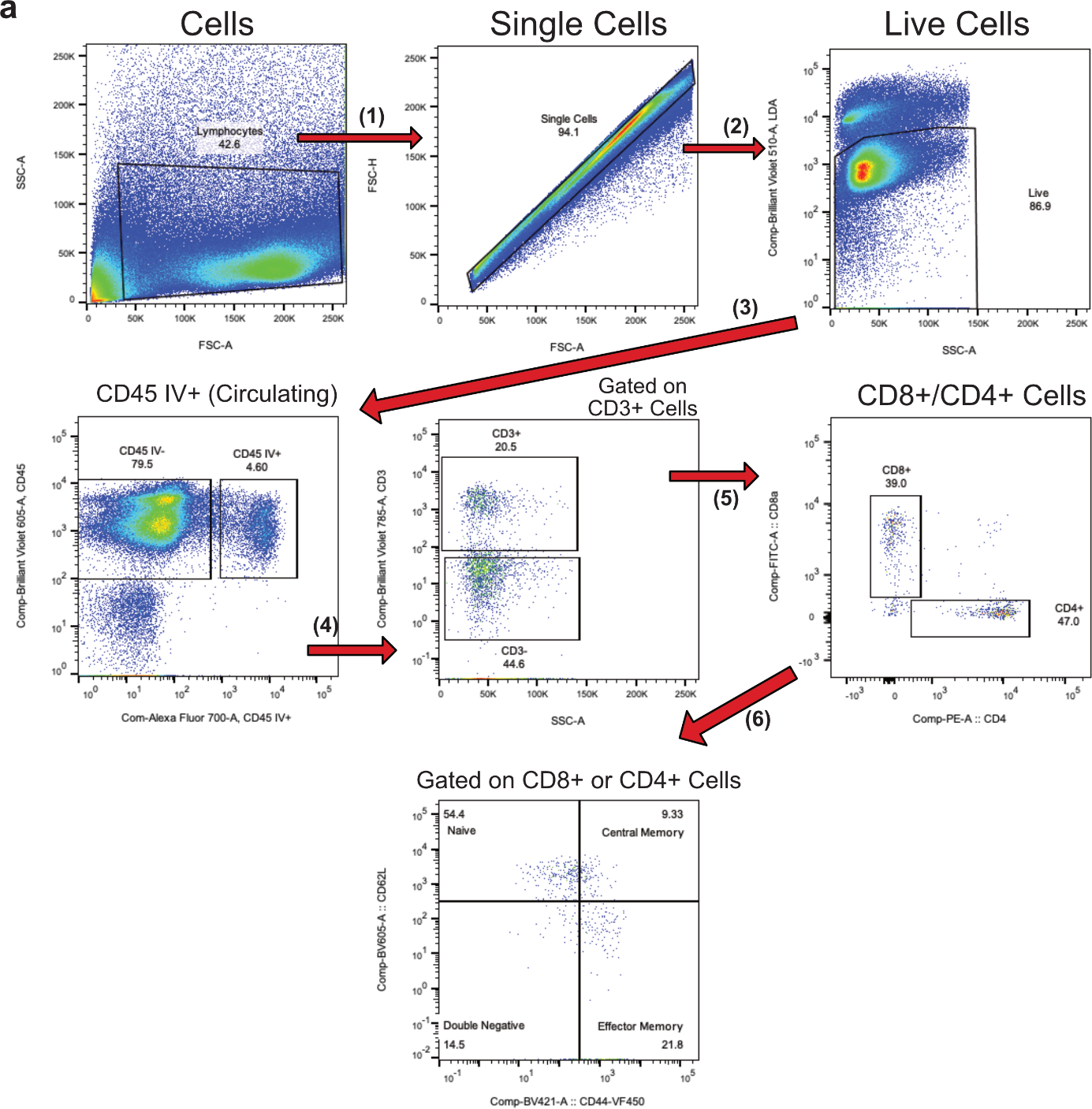

**Supplemental Figure 5. Assessment of circulating T-cell surface phenotype in spleen and lungs upon distinct diet regimens in male mice post-vaccination and post-influenza challenge.** (a) Gating strategy for identifying live CD45 IV^+^ CD3^+^ cells in mouse spleen and lung samples. (b) Each flow plot represents cells gated on live CD45 IV^+^ CD3^+^ from an individual sample as indicated in the gating strategy. Naïve (CD62L^+^CD44^-^), Central Memory (T_CM_, CD62L^+^CD44^+^), and Effector Memory (T_EFM_, CD62L^-^CD44^+^) cells are labeled on representative flow plots.

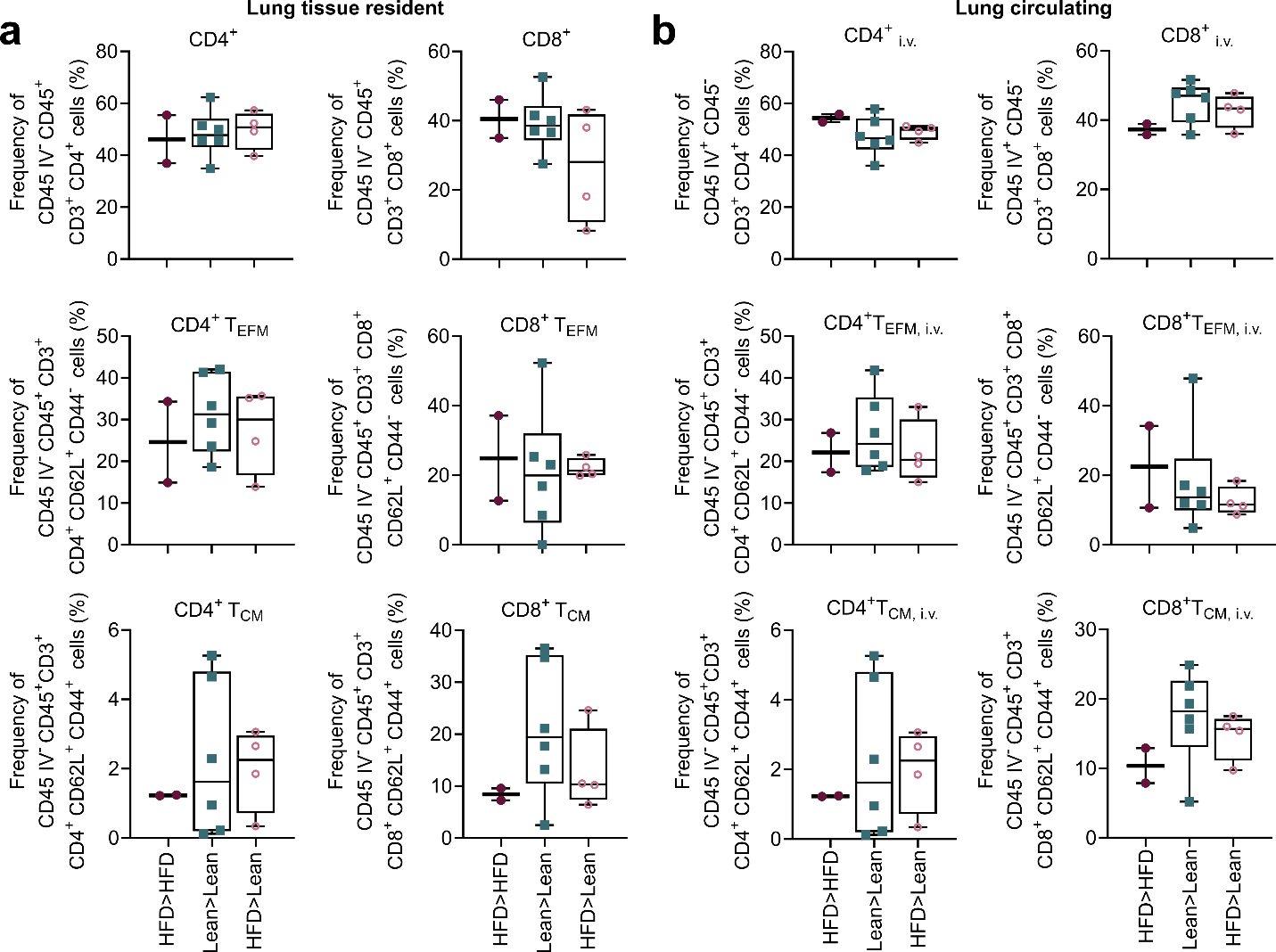

**Supplementary Figure 6. Frequency plots of recall response to vaccination before diet switch.** Lungs were harvested and homogenized into single-cell suspensions and 1 × 10^6^ cells were stained for CD4^+^ T cells, CD8^+^ T cells, and B220^+^ B cells at 7 days post-challenge via flow-cytometry. (a) Frequency of lung resident CD45_i.v.−_, CD4^+^ and CD45_i.v.−_, CD8^+^ T cells; CD45_i.v.−_, CD4^+^ effector memory (T_EFM_) and CD45_i.v.−_, CD8^+^ T_EFM_ cells; CD45_i.v.−_, CD4^+^ central memory (T_CM_) and CD45_i.v.−_, CD8+ T_CM_ cells. (b) Frequency of circulating CD45_i.v.−_, CD4^+^ and CD45_i.v.−_, CD8^+^ T cells; CD45_i.v.−_, CD4^+^ effector memory (T_EFM_) and CD45_i.v.−_, CD8^+^ T_EFM_ cells; CD45_i.v.−_, CD4^+^ central memory (T_CM_) and CD45_i.v.−_, CD8+ T_CM_ cells. Data in include always obese (HFD>HFD), always lean (Lean>Lean), and formerly obese (HFD>Lean) mice (n = 2, n=6, n=4 for each group, respectively). Data were analyzed using FlowJo version 10.8.1. Outliers in the dataset were removed using the robust regression and outlier removal (ROUT) method (Q=1%). Graphed in as means ± standard error with comparisons made between diet groups via paired T-test. Red closed circles = always obese mice, green closed squares = always lean mice, and pink open circles = formerly obese mice.

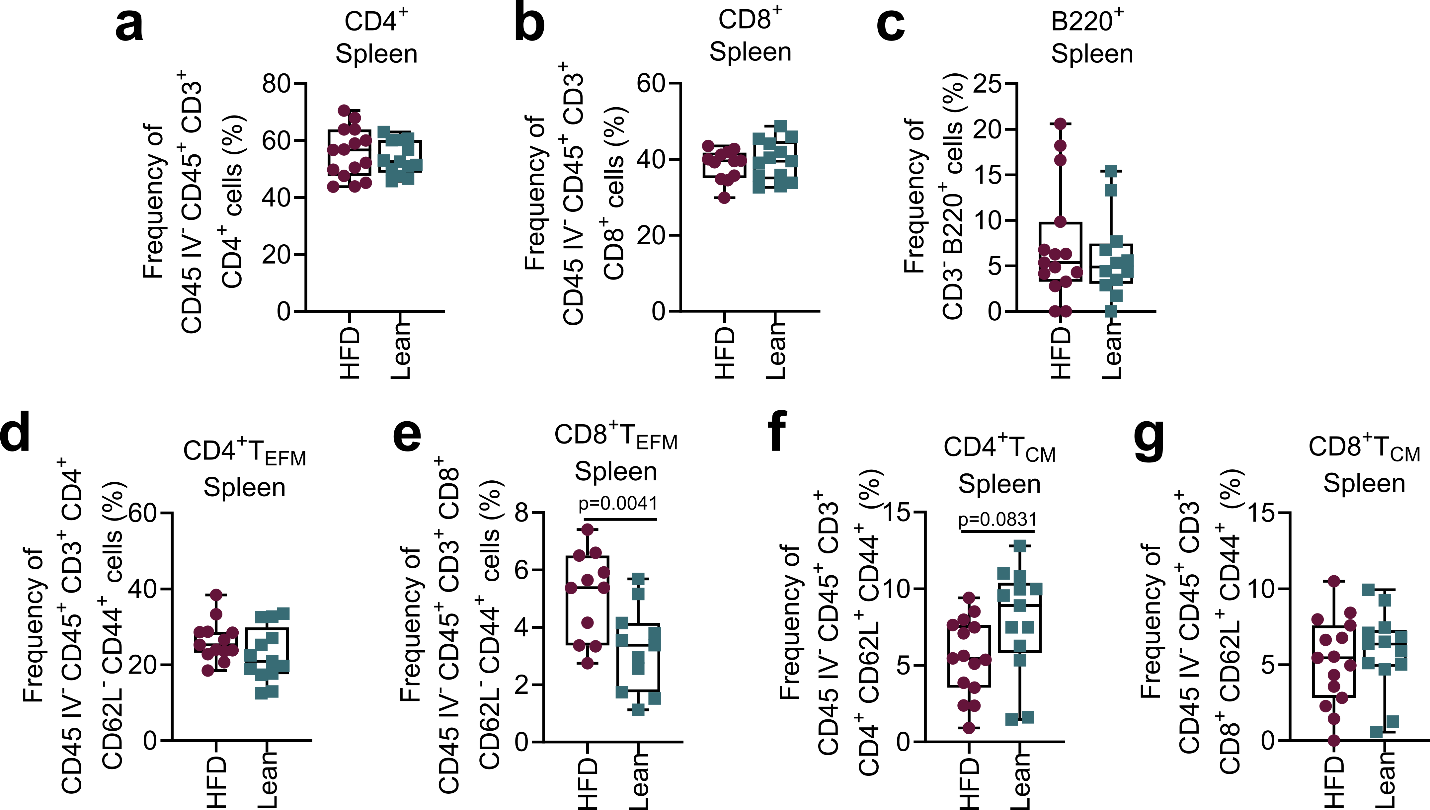

**Supplemental Figure 7. Frequency plots of primary response to vaccination before diet switch.** Spleens were harvested and homogenized into single-cell suspensions and 1 × 10^6^ cells were stained for CD4^+^ T cells, CD8^+^ T cells, and B220^+^ B cells at 21 days post-vaccination via flow-cytometry. (a-g) Frequency of (a) CD4^+^ and (b) CD8^+^ T cells and (c) B cells, (d) CD4^+^ effector memory (T_EFM_) and (e) CD8^+^ T_EFM_ cells, (f) CD4^+^ central memory (T_CM_) and (g) CD8+ T_CM_ cells. Note that data in include always obese/future formerly obese group (HFD) and always lean (Lean) mice (n = 16 per group) as samples were collected prior to diet switch. Data were analyzed using FlowJo version 10.8.1. Outliers in the dataset were removed using the robust regression and outlier removal (ROUT) method (Q=1%). Graphed in as means ± standard error with comparisons made between diet groups via paired T-test. Red closed circles = always obese mice, green closed squares = always lean mice, and pink open circles = formerly obese mice.

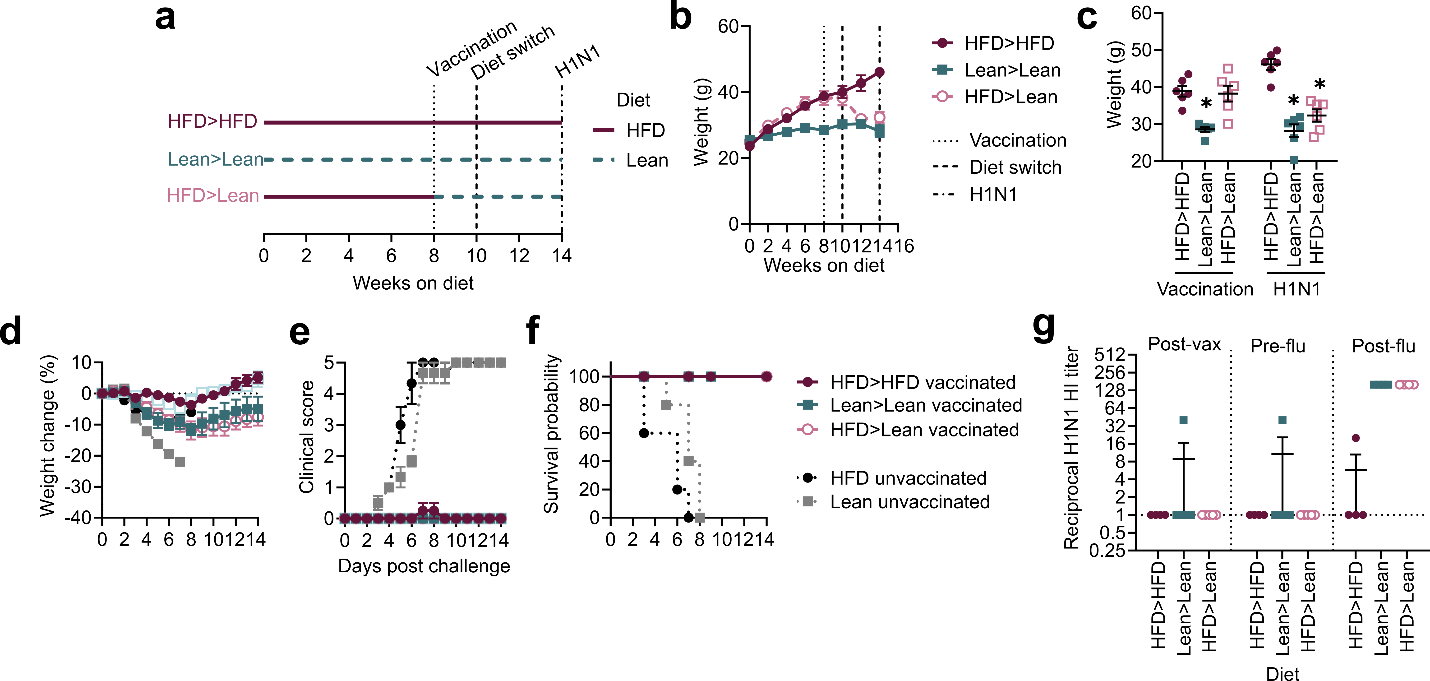

**Supplemental Figure 8. Increased survival in phenotypic obese and formerly obese mice with 8-week HFD exposure.** (a) Timeline of diet administration, vaccination, and challenge in half-term diet switch studies. (b) Weights of mice through a half-term diet switch studies (n=5-10 mice/group) at (c) vaccination and challenge. (d-f) Morbidity and mortality upon H1N1 virus challenge with n=5-10 mice/group. (d) Weight curves with statistical comparisons made via mixed-effects model, (e) clinical scores with statistical comparisons made using a two-way ANOVA, and (f) survival with statistical comparisons made using Mantel-Cox log-rank analysis. (g) Hemagglutination inhibition titers at indicated timepoints. Data are represented in (b-e) as means ± standard error, in (f) as surviving proportions with censored or event animals indicated by their respective symbols, and in (g) as geometric mean ± standard deviation with x indicating no mice surviving to time point. Statistical comparisons test between HFD>HFD and indicated diet groups (n=10 mice/group) in (b, f) and between mock and vaccinated animals within a single diet group in (c-e). Red closed circles = always obese mice, green closed squares = always lean mice, pink open circles = formerly obese mice, black closed circle = mock vaccinated obese mice, grey closed square = mock vaccinated lean mice.

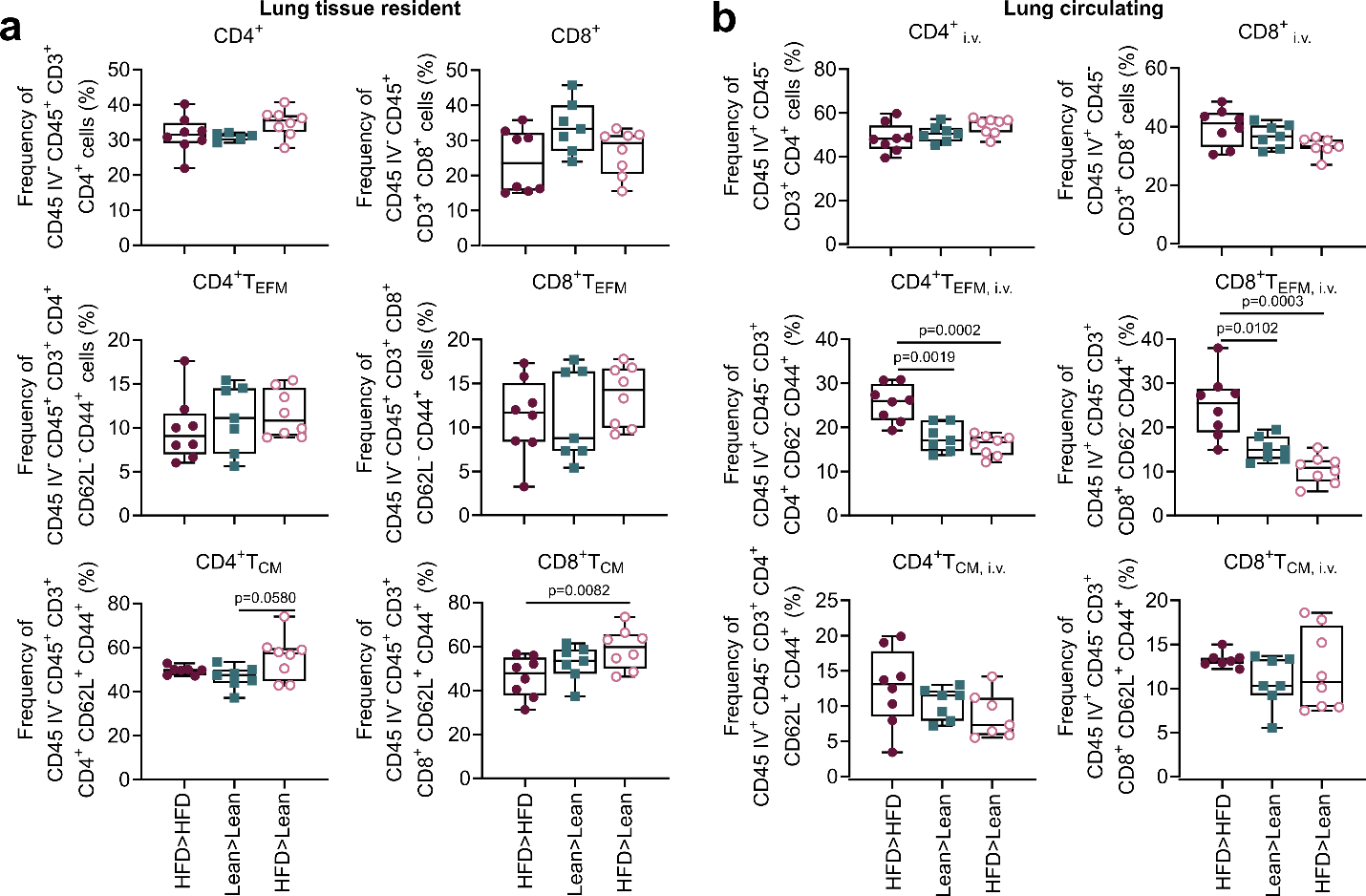

**Supplemental Figure 9. Frequency plots in lungs of diet-switched mice before vaccination and at 7 days post-challenge.** Lungs were harvested and homogenized into single-cell suspensions and 1 × 10^6^ cells were stained for CD4^+^ and CD8^+^ T cells at 7 days post-challenge and quantified via flow-cytometry. (a) Frequency of lung resident CD45_i.v.−_, CD4^+^ and CD45_i.v.−_, CD8^+^ T cells; CD45_i.v.−_, CD4^+^ effector memory (T_EFM_) and CD45_i.v.−_, CD8^+^ T_EFM_ cells; CD45_i.v.−_, CD4^+^ central memory (T_CM_) and CD45_i.v.−_, CD8+ T_CM_ cells. (b) Frequency of circulating CD45_i.v.+_, CD4^+^ and CD45_i.v.+_, CD8^+^ T cells; CD45_i.v.+_, CD4^+^ T_EFM_ and CD45_i.v.+_, CD8^+^ T_EFM_ cells; CD45_i.v.+_, CD4^+^ T_CM_ and CD45_i.v.+_, CD8+ T_CM_ cells. Data were analyzed using FlowJo version 10.8.1. Outliers in the dataset were removed using the robust regression and outlier removal (ROUT) method (Q=1%). Data include always obese (HFD>HFD), always lean (Lean>Lean), and formerly obese (HFD>Lean) mice (n = 8, n=7, n=7 for each group, respectively) and graphed as means ± standard error with comparisons made between diet groups via two-way ANOVA. Red closed circles = always obese mice, green closed squares = always lean mice, and pink open circles = formerly obese mice.
